# A modular cloning toolbox including CRISPRi for the engineering of the human fungal pathogen and biotechnology host *Candida glabrata*

**DOI:** 10.1101/2022.10.17.512482

**Authors:** Sonja Billerbeck, Rianne C Prins, Malte Marquardt

## Abstract

The yeast *Candida glabrata* is an emerging, often drug-resistant opportunistic human pathogen, that can cause severe systemic infections in immunocompromised individuals. At the same time, it is a valuable biotechnology host that naturally accumulates high levels of pyruvate – a valuable chemical precursor. Tools for the facile engineering of this yeast could greatly accelerate studies on its pathogenicity and its optimization for biotechnology. While a few tools for plasmid-based expression and CRISPR/Cas-based genome engineering have been developed, there is no well-characterized cloning toolkit available that would allow the assembly of pathways or genetic circuits in a modular fashion. Here, by re-using and characterizing the *Saccharomyces cerevisiae*-based yeast molecular cloning toolkit (YTK) in *C. glabrata* and by adding missing components, we build a well-characterized CgTK (*Candida glabrata* toolkit). We used the CgTK to build a CRISPR interference system for *C. glabrata* that can be used to generate selectable phenotypes via single-gRNA targeting such as required for genome-wide library screens.

## INTRODUCTION

*Candida glabrata* is an opportunistic human fungal pathogen, that causes 10-25% of fungal bloodstream infections in humans.^1–5^ Clinical isolates of *C. glabrata* are increasingly resistant to multiple drugs and as a consequence, mortality rates of blood-stream infected individuals are high as no appropriate treatment is available.^6–9^ Unlike other human fungal pathogens that are acquired from the environment, *C. glabrata* is a natural commensal of the human mycobiome, mostly the GI tract,^1,10,11^ but it can overgrow and turn virulent once the immune system of the human host is compromised. The virulent phenotype is linked to *C. glabrata’s* ability to grow rapidly at 37°C, its high capacity for adhesion – based on the presence of a large repertoire of adhesin genes – and biofilm formation, its ability to intrinsically tolerate certain antifungal drugs, and its rapid adaptation to stresses.^8,12–14^ For example, it can sustain prolonged starvation periods, tolerate oxidative stress and survive or even thrive within the acidic environment of phagosomes.^15–18^ These traits are activated by various environmental cues, such as pH, carbon source, and temperature.^19^ Although much progress has been made in understanding *C. glabrata’s* biology, the molecular regulatory underpinnings that enable the phenotypic adaptation to a pathogenic lifestyle are not yet understood.

At the same time, some *C. glabrata* strains are used as valuable chassis for biotechnology as they naturally produce high levels of pyruvic acid when grown under high glycolytic flux conditions.^20,21^ Pyruvate is an important precursor for several chemical synthesis processes and agro-chemistry, is directly useful as a dietary supplement and is used in pharmaceutical applications. In addition, *C. glabrata* has been engineered to metabolize pyruvic acid into valuable downstream products such as α-ketoglutarate,^22^ fumarate,^23^ acetoin,^24^ malate^25^ and diacetyl.^26^

Tools enabling a synthetic biology approach (based on design-build-test-learn cycles) to *C. glabrata’s* biology could not only facilitate its metabolic engineering towards biotechnological applications but also help elucidate the genetic and regulatory architectures underlying the phenotypic transition to virulence in this opportunistic pathogen.^27^ In addition, understanding the genetic design behind *C. glabrata’s* phenotypic adaptation towards virulence could become a valuable inspiration for Synthetic Biology and help design sophisticated programmed cellular behavior.

One step towards facilitating synthetic biology projects in *C. glabrata* is the development of a toolkit with well-characterized genetic parts and gene regulation tools for synthetic circuit design. Modular cloning kits based on Golden Gate assembly^28^ have proven indispensable to overcome cloning barriers for bacterial and eukaryotic hosts, including plants.^29–32^ They provide standardized and well-characterized parts that enable the rapid assembly of genetic circuits and pathways, reduce the number of re-design cycles and thus allow for a faster scientific outcome. This is relevant as cloning is often the gate-opener to answering advanced biological questions but often remains the temporal bottleneck. As such, modular cloning kits lower the burden of technical cloning details, allowing to focus on higher-level experimental design.

Some tools already exist for *C. glabrata*, such as a set of expression plasmids,^33^ several non-homologous-end-joining-based CRISPR tools^34,35^ as well as transposons for gene disruption,^36,37^ and a deletion collection covering 12% of the *C. glabrata* genome.^38^ Here, we extend the genetic toolbox for *C. glabrata* by building a molecular cloning toolkit (CgTK) by re-characterizing and extending the Yeast Toolkit (YTK) built for *Saccharomyces cerevisiae*. Despite its “Candida” clade name, *Candida glabrata* is evolutionarily closely related to *S. cerevisiae*^12^ and several *S. cerevisiae*-derived promoters and auxotrophic markers have shown to be functional in *Candida glabrata*.^33^ Still lacking is the comprehensive characterization of gene expression units that could allow for predictable circuit design or metabolic engineering in *C. glabrata*. Eventually, we use the CgTK to develop a CRISPR interference (CRISPRi) system for single-gRNA expression knock-down. Currently, there are no CRISPRi systems available for *C. glabrata*. On the way, we also show the first experimental verification that the *C. glabrata* open reading frame CAGL0A04587g is indeed a functional ortholog of the *ALG3* gene in *S. cerevisiae* – in *S. cerevisiae* encoding for an alpha-1,3-mannosyltransferase involved in (membrane)-protein glycosylation – as its deletion or repression renders *C. glabrata* resistant to the killer toxin HMK/HM-1.

## RESULTS AND DISCUSSION

### Design of the *Candida glabrata* toolkit (CgTK)

Given the close relationship between *S. cerevisiae* and *C. glabrata*,^12^ we reasoned that the most resource-saving way to build the CgTK was to re-use the *S. cerevisiae* YTK and characterize part performance in *C. glabrata* such that these data are available for experimental design. Further, reusing the *S. cerevisiae* parts gives a large repertoire of parts that are sequence-orthogonal to the *C. glabrata* genome. Besides re-using existing parts, we identified missing and non-performing parts and added or replaced these parts to make the CgTK most useful (**Figure 1** and **Supplementary Table 1 and 2)**. In brief, in total, we characterized 21 constitutive yeast promoters (19 YTK, 2 new), four constitutive synthetic minimal promoters (4 new), three inducible promoters (1 YTK, 2 new), and three protein degradation tags (3 YTK). We constructed nine vector sets for the assembly of (multiple) transcriptional units featuring four auxotrophic markers and two origins of replication. Venus and mRuby2 were used as read-outs for protein expression levels from different constructs and fluorescence was calibrated using previously described calibrant dyes for green and red fluorescence.^39^

**Figure 1:**
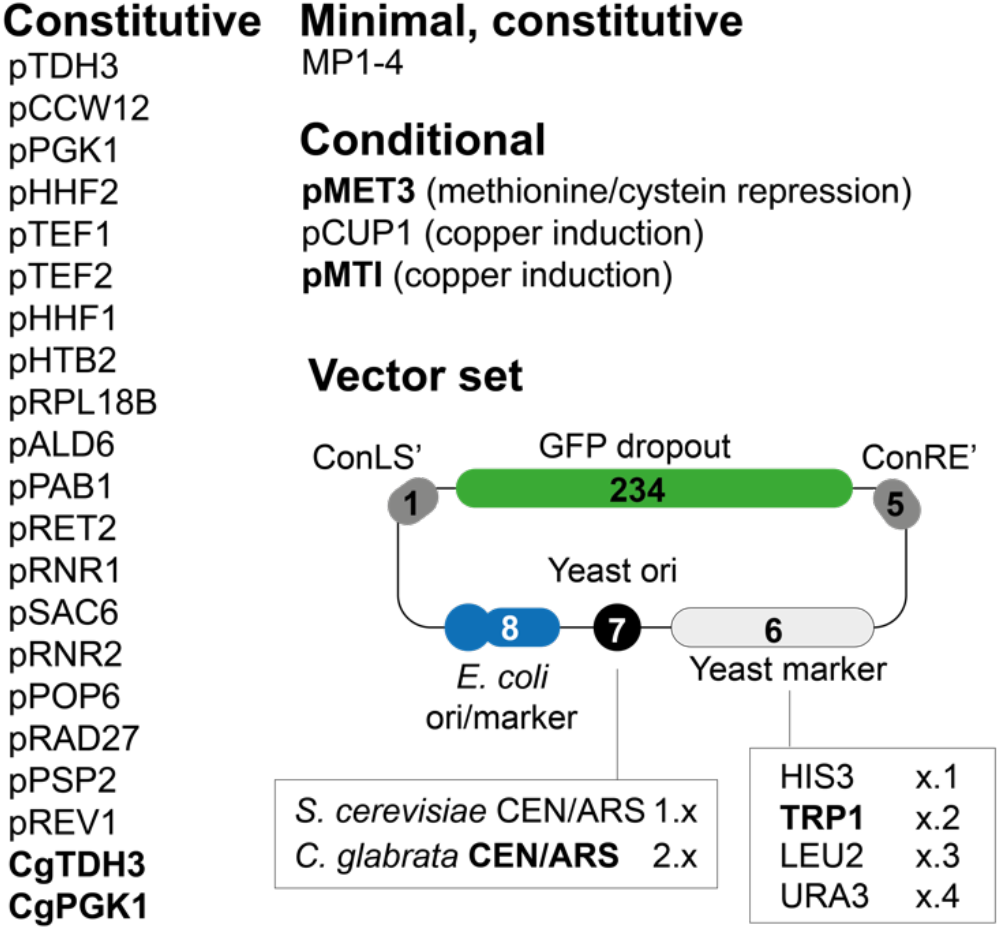
Overview on the CgTK. Overview on the YTK-parts that were characterized in *C. glabrata* and newly added parts (indicated in bolt). In total, the CgTK contains 26 constitutive promoters (19 YTK, 2 from *C. glabrata* and 4 minimal promoters), three inducible/repressible promoters (Copper and methionine/cysteine), three degradation tags, and nine pre-assembled vectors with different selection markers (eight auxotrophic vectors as shown here and one with a nourseotricin marker, **Supplementary Table 2**)

All experiments were performed in *C. glabrata* ATCC 2001 HTL^−^.^38^ This strain is a clinical isolate frequently used for *C. glabrata* research including the creation of the *C. glabrata* deletion collection ^38^. *C. glabrata* ATCC 2001 HTL^−^ is auxotrophic for histidine biosynthesis (*HIS3*), leucine biosynthesis (*LEU2*), and tryptophan biosynthesis *(TRP1)* (*his3*Δ::FRT *leu2*Δ::FRT *trp1*Δ::FRT). In addition, we created a variant auxotrophic for uracil biosynthesis (U^−^) via a simple 5-FOA selection protocol.^40^

### A set of cloning vectors with *S. cerevisiae* (Sc) and *C. glabrata* (Cg) CEN/ARS origins of replication and four auxotrophic markers

First, to facilitate the cloning of single and multiple transcriptional units based on the YTK-Golden Gate scheme, we created and tested an eight-membered vector set featuring the YTK-derived HIS3 (pYTK76), LEU2 (pYTK75) and URA3 (pYTK74) selection markers and a herein-cloned TRP1 selection marker (pCgTK02, TRP1 transcriptional unit from *C. glabrata*) **(Figure 1)**. All selection markers were combined with either the YTK*-*derived CEN/ARS origin of replication from *S. cerevisiae* (pYTK81) or a herein created *C. glabrata* derived CgCEN/ARS origin of replication (pCgTK01). Most *C. glabrata* research relies on plasmids coding for the CgCEN/ARS while most available yeast plasmids (e.g. on Addgene) are based on an *S. cerevisiae*-derived CEN/ARS replication origin. To facilitate the use of plasmids designed for *S. cerevisiae* in *C. glabrata*, we were interested in whether the ScCEN/ARS sequence is functional in *C. glabrata* and if there would be a difference in performance when compared to the CgCEN/ARS. Further, we equipped the vector set with the GFP dropout cassette (pYTK47) flanked by the ConLS’ and ConLR’ connectors (pYTK08, pYTK73) for single and multiple TU cloning followed by green/white screening in *Escherichia coli*.

*C. glabrata* cells were transformed with all vectors and the recovery of transformants on all corresponding selective plates indicated that all auxotrophic markers were able to functionally complement the strain’s metabolic auxotrophies and that all origins of replication were functional. Next, we recorded growth curves to detect potential differences in complementation capacity or plasmid burden. Vector-specific plasmid burden had been shown before for *S. cerevisiae* and can help in choosing the right vector for an application.^41^

We measured the growth of strains transformed with the “empty” vectors (vectors encoding a GFP-dropout as 2/3/4 part, see Lee *et al*.^29^ for part descriptions within the YTK) and vectors carrying an HHF1p-Venus-ENO1t transcriptional unit as 2/3/4 part **(Supplementary Figures 1A and 2A)**. Cells carrying the empty vector showed overall similar growth behavior on their respective selective media when compared to cells not carrying a vector on non-selective media **(Supplementary Figures 1B and C)**. Only cells carrying vector version 1.3 (ScCEN/ARS, LEU2) showed slightly slowed growth and reduced final OD_630_ **(Supplementary Figure 1B)**. The increased impact of the *LEU2* marker on growth in *C. glabrata* is consistent with results using this marker in *S. cerevisiae*.^41^

In contrast to the empty vectors, cells carrying the vector set encoding the HHF1p-Venus-ENO1t transcriptional unit showed more pronounced growth phenotypes: while the CgCEN/ARS vectors showed very minor growth differences **(Supplementary Figure 2C)**, the ScCEN/ARS showed longer lag phases and reached only 82% (version 1.1, ScCEN/ARS, HIS3), 66% (version 1.2, ScCEN/ARS, TRP1) and 59% (version 1.3, ScCEN/ARS, LEU2) of the final OD_630_ of the untransformed control **(Supplementary Figure 2B)**.

### Copy number differences across the vector set allow for tuning expression strength

Given the growth phenotypes, we were interested in vector-dependent differences in the expression levels and potential differences in plasmid copy number. We first tested expression levels using the vectors containing the HHF1p-Venus-ENO1t transcriptional unit. The ScCEN/ARS-based vectors showed a 3 to 8-fold higher expression level when compared to their CgCEN/ARS-based counterparts with the same auxotrophic marker **(Supplementary Figure 3)**. The highest expression was observed for the vector version 1.1 (ScCEN/ARS, HIS3). This vector showed a 3-fold higher expression level than the other ScCEN/ARS-based vectors. Version 1.2 (ScCEN/ARS, TRP1) and version 1.3 showed similar expression levels when compared to each other. Finally, we cross-compared the relative copy numbers of some of the vectors by qPCR **(Supplementary Figure 4)**: we used two different DNA extraction methods, one for complete DNA extraction and one for plasmid extraction, and measured the abundance of the Venus gene in cells carrying vector version 1.1, 1.2, 2.1 and 2.2. Both methods showed that vector version 1.1 (ScCEN/ARS, HIS3) - which had shown the highest expression level - was the most abundant vector, while vector version 2.2 (CgCEN/ARS, HIS3) was the least abundant. The plasmid extraction showed that vector 1.2 and 2.1 were 4-fold less abundant than vector version 1.1. Vector version 2.2 was 29 times less abundant than vector 1.1 **(Supplementary Figure 4A)**.

In summary, our data point to significant differences in copy number across the tested vectors and point to the fact that not only the origin of replication but also the selection marker seems to impact copy number. Further, the data show that copy number alone is not responsible for growth burden or expression levels: For example, vector 2.1 (CgCEN/ARS, HIS3) seems to be as abundant as vector 1.2 (ScCEN/ARS, TRP1) but it shows less growth burden **(Supplementary Figure 2)** but also lower expression from the HHF1 promoter **(Supplementary Figure 3)**. In line with similar plasmid-growth-burden-related findings for *S. cerevisiae*, this indicates the need for good experimental characterization of parts to allow predictable engineering.^41^

### The high copy 2μ origin of replication from *S. cerevisiae* is not functional in *C. glabrata*

While the above characterized Sc/CgCEN/ARS system delivered a vector set with different expression strength likely due to difference in copy number, we were further interested if the 2μ high-copy origin of replication was functional in *C. glabrata*, but we were not able to recover transformants with any selection marker indicating that this origin does not replicate in *C. glabrata*.

### The nineteen YTK*-*derived constitutive promoters are functional in *C. glabrata*

Next, we characterized the performance of all 19 YTK-derived constitutive promoters in *C. glabrata* using two different vector backbones; the highest expression vector 1.1 (ScCEN/ARS, HIS3) and one of the low expression vectors, vector version 2.2 (CgCEN/ARS, TRP1). We used the Venus fluorescent reporter (pYTK33) combined with the YTK-derived ENO1 terminator (pYTK51) as readout.

Promoters were characterized by measuring bulk fluorescence in a plate reader using a very similar protocol as the original YTK study.^29^ Arbitrary fluorescence units were then normalized to the chemical calibrant fluorescein to facilitate lab-to-lab portability of the data.^39^ **Figures 2A** and **B** show that all 19 YTK promoters were functional in *C. glabrata*, showing at least 3-fold expression over background in the higher expression vector version 1.1 **(Figure 2A)**. The background was defined as the autofluorescence of *C. glabrata* cells not harboring a plasmid. In line with our previous result, expression from vector version 1.1 yielded higher expression across promoters when compared to vector version 2.2 **(Figures 2A and B)**. The fold-difference in expression between vector 1.1 and 2.2 was promoter-dependent and varied from 8-fold to 30-fold for the six strongest promoters **(Supplementary Figure 5)**. Although the exact order of strongest-to-weakest promoters slightly varied between vector version 1.1 and 1.2 we could identify a set of strong, medium, and weak promoters **(Figure 2C)**. Overall the 19 promoters covered two orders of magnitude in expression strength when expressed from vector version 1.1 **(Figure 2D)**. We also compared the expression strength of all promoters in vector version 1.1 to expression levels in *S. cerevisiae* (BY4741). Interestingly, several of the promoters showed different strength profiles in *C. glabrata* when compared to *S. cerevisiae* **(Figure 2D)**. For example, the TDH3 and HHF2 promoters ranked among the strongest promoters in *S. cerevisiae* while they showed medium (TDH3p) to low (HHF2p) expression in *C. glabrata* (**(Figure 2D)**. On the other hand, the PAB1 promoter showed medium strength in *S. cerevisiae* and is a rather strong promoter in *C. glabrata*.

**Figure 2.**
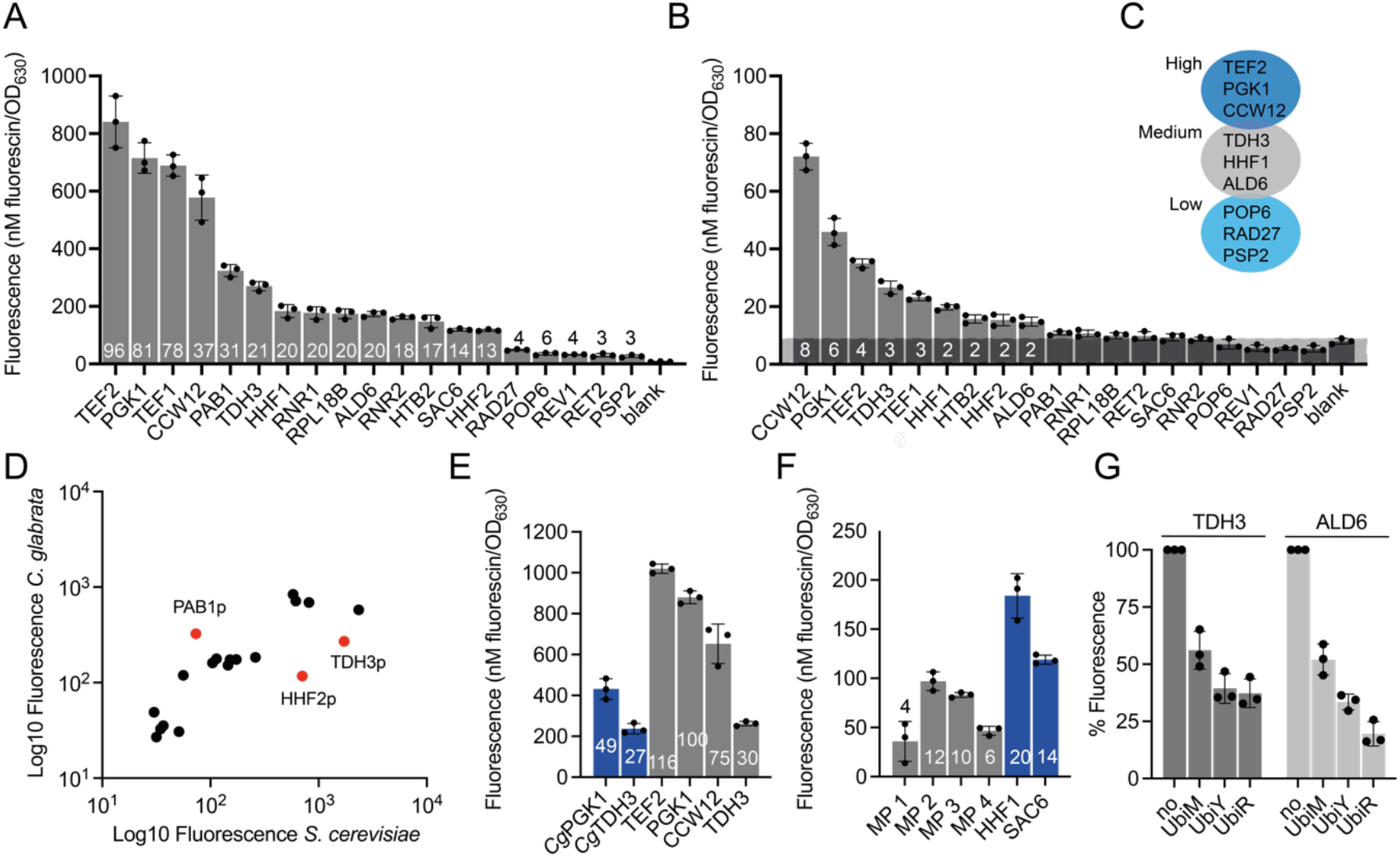
Performance of YTK and additional parts in *C. glabrata* using green fluorescence (Venus) as readout. **A and B:** Performance of the 19 YTK promoters in *C. glabrata* using two different vectors; version 1.1 (A) and 2.2 (B). The numbers indicate the fold-change in fluorescence over background, where background is defined as the autofluorescence of *C. glabrata* cells not carrying a plasmid. Note: The Y-axis shows a different scale. **C:** Suggested high, medium and low expression promoters based on the data displayed in A and B. **D:** Comparison of promoter performance in *C. glabrata* and *S. cerevisiae*. The data that was generated with vector version 1.1 is used. Outliers are marked in red. **E:** Comparison of *C. glabrata* derived TDH3 and PGK1 promoters with the strongest YTK-derived promoters. Data were generated in vector version 1.1 using green fluorescence of the Venus protein as readout. **F:** Performance of minimal yeast promoters in *C. glabrata*, promoters are compared to the YTK-derived HHF1p and SAC6p. **G:** Performance of the protein degradation tags Ubi-M, Ubi-Y, and Ubi-R in *C. glabrata*. All experiments were run in biological triplicates (three transformants) and error bars represent the standard deviation.

Absolute achievable expression strength within the set of promoters was 2.7-fold lower in *C. glabrata* when compared to *S. cerevisiae* **(Supplementary Figure 6)**. To test if we could reach higher expression strength in *C. glabrata* we cloned two additional *C. glabrata*-endogenous promoters from the central carbon metabolism, the CgTDH3 promoter and the CgPGK1 **(Figure 2E)**. Neither promoter showed a higher expression than the YTK-promoters, but both promoters could be used as medium to strong promoters with 49-fold and 27-fold expression over background.

We further tested the performance of a subset of seven promoters with a red fluorescent protein (mRuby2, YTK034) as readout. All arbitrary fluorescence values were normalized to the calibrant dye sulforhodamine-101 **(Supplementary Figure 7)**.^39^ Similar to the green fluorescence, also for the red fluorescence vector version 1.1 showed higher expression than 2.2 (between 11 and 2-fold difference depending on promoter **(Supplementary Figure 7A, B and D)**. Vector version 1.1 yielded a 44-fold expression over background with the strongest promoter tested (the medium-level TDH3 promoter) and vector version 2.2 a 4-fold expression level over background for the same promoter.

Overall, mRuby expression yielded about 10-fold less fluorescence when compared to Venus expression, measured and compared in nM calibrant dye. The lower fluorescence output with mRuby2, when compared to Venus, was also observed in *S. cerevisiae* BY4741, both in the original YTK data set^29^ and in our data set **(Supplementary Figure 7C)**.

### Minimal *S. cerevisiae* synthetic promoters are functional in *C. glabrata*

We further cloned and tested four constitutive minimal synthetic promoters (MPs) that were originally engineered for performance in *S. cerevisiae*.^42^ The advantages of these promoters is their short length (∼129 bp versus 700 bp) and modularity. Various core promoters of 30 bp can be combined with constitutive or inducible upstream activating sequences (UAS) of 10 bp each, to create promoters of various strengths.^42^ Here we used the core promoters 1, 5, 8 and 9, which had shown different strengths in *S. cerevisiae*. We combined these with the UASs E, F, and C to design the minimal promoters MP1 to MP4. All four promoters were functional in *C. glabrata* yielding between 4 times (MP1, low expression) and 12 times (MP2, medium expression) expression over background. None of the promoters yielded strong expression, but also expression levels in *S. cerevisiae* were medium for all the promoters **(Supplementary Figure 6C)**. It has been shown before that the YTK Golden Gate design leaves a BglII seam between the promoter and the ORF which is suboptimal for expression strength and can be improved by re-coding.^43^ This might explain why the MP design did not achieve the reported high expression levels in this specific set-up.

In conclusion, the minimal promoters (MPs) can be useful as short low to medium expression promoters in *C. glabrata* and they could serve as a starting point to further optimize their design to yield higher expression within the YTK Golden Gate design.

### Three inducible promoters for the use in *C. glabrata*

The YTK contains two inducible promoters: 1) the galactose inducible promoter GAL1p and 2) the copper (II) sulfate inducible promoter CUP1p. It is known that the *S. cerevisiae*-derived galactose inducible promoter is not functional in *C. glabrata*, as it lacks the molecular machinery for galactose metabolism and transport, including the galactose-specific transcriptional activator GAL4. Copper induction with the *S. cerevisiae* CUP1 promoter has not been tested to the best of our knowledge.

To add inducible parts, we cloned and characterized two additional inducible *C. glabrata* promoters: the methionine and/or cysteine repressible promoter MET3p and the *C. glabrata* copper (II) sulfate inducible promoter MT-1p. In analogy to the constitutive promoter testing, the three inducible promoters, MET3p, CUP1p, and MT-1p were tested in the high- and low-expression vectors version 1.1 and 2.2.

Figure 3 summarizes the performance of the MET3 promoter when repressed with cysteine or methionine. Cells were grown in media without methionine (a usual component of minimal drop-out media in yeast) and 2-fold dilutions starting from 10 mM methionine or cysteine were externally added. The MET3 promoter in vector version 1.1 allowed for a 17-fold change in Venus expression when using methionine as a repressor. Full repression was achieved at a concentration of ≧625 μM methionine and remaining promoter leakiness were 2-fold over background **(Figure 3A and C)**. When using cysteine as a repressor, a 7-fold change in expression was achieved. The reduction in fold change was mainly due to the fact that cysteine was less effective at repressing the promoter. At full cysteine repression (≧5 mM cysteine), leakiness was still 3-fold over background. Interestingly, using cysteine as a repressor widened the linear range of the promoter when compared to methionine (Figure 3A), suggesting that - depending on the application - full repression can be achieved with methionine, while medium levels could be finer tuned with cysteine. Of note, 10 mM cysteine inhibited the growth of *C. glabrata* **(Supplementary Figure 8)**, as such only data for 5 mM cysteine were included **Figure 3A**. Similar results were achieved when using vector version 2.2, but in line with previous results, overall expression levels were reduced **(Figure 3B and D)**. Methionine repression allowed for a 4-fold change in expression level, leaving no detectable background expression at a concentration of ≧625 μM methionine. Cysteine repression allowed for a 3-fold change with no detectable background expression at concentration of ≧2.5 mM cysteine. In analogy to the other vector, cysteine resulted in a wider dynamic range of the promoter, leaving the opportunity to use a combination of methionine (for full repression) and cysteine (tuning) to achieve the desired expression level. The MET3 promoter from *C. glabrata* was not functional in *S. cerevisiae*.

**Figure 3.**
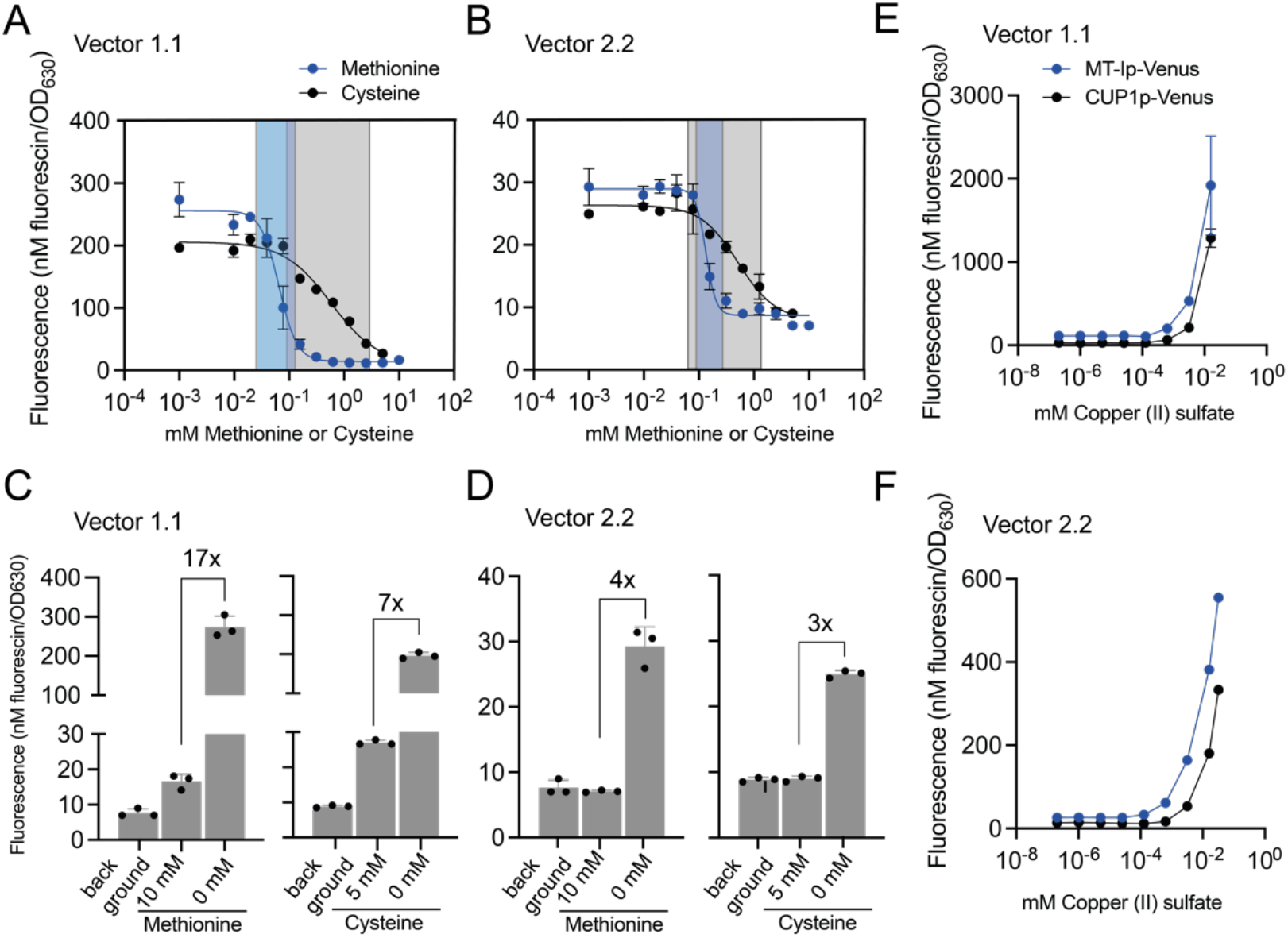
Performance of the methionine/cysteine repressible and copper inducible promoters. **A-B:** Performance of MET3 promoters in vector version 1.1 (A) and 2.2 (B) when grown in the presence of decreasing concentrations of methionine (starting from 10 mM methionine and 5 mM cysteine, two-fold dilutions were measured). **Note:** 10 mM cysteine led to partial growth inhibition **(Supplementary Figure 9)**, this is why the 5 mM value is displayed. The grey and blue shaded boxes indicate the dynamic range of the promoter repressed methionine (blue) and repressed with cysteine (grey). **C-D:** Fold-change difference in expression level between the “ON” and the “OFF” state of the MET3 promoter in vector version 1.1 (C) and 2.2 (D) when induced with 10 mM methionine or 5 mM cysteine. The “OFF” state is also compared to the background autofluorescence of cells without a plasmid (blank). **E-F:** Performance of the MT-I and the CUP1 promoters in vector version 1.1 (E) and 2.2 (F) when grown in the presence of increasing concentrations of copper (II) sulfate (starting from 2 mM methionine, five-fold dilutions were measured). Note: *C. glabrata’s* growth is (partly) inhibited by copper concentration above 16 μM **(Supplementary Figure 11)**.

**Figure 3 E-F** shows the performance of the two copper (II) inducible promoters CUP1p and MT-1p in *C. glabrata*. For comparison, we also characterized both promoters in *S. cerevisiae* **(Supplementary Figure 9)**. First, we noticed that *C. glabrata* cells were much more sensitive to copper (II) sulfate (CuSO_4_) than *S. cerevisiae* cells **(Supplementary Figure 10**). For induction, we used 5-fold dilutions starting from 2 mM copper (II) sulfate and *C. glabrata* cells would only grow at concentrations lower than 16 μM (though reaching lower final OD_630_) and would only reach full OD_630_ at concentrations lower than 3 μM. In contrast, *S. cerevisiae* cells reached full OD_630_ at the highest tested concentration of 2 mM copper (II) sulfate. Still, *C. glabrata* cells seemed metabolically active and produced high levels of Venus protein at concentrations above 16 μM. Interestingly, in analogy to the MET3 promoter, the MT-1 promoter was not functional in *S. cerevisiae*, not leading to any expression above background in neither presence nor absence of inducer **(Supplementary Figure 9)**.

### Porting the CgTK into clinical isolates of *C. glabrata*

As the MET3 promoter seemed to be a very suitable inducible and tunable promoter for *C. glabrata*, we were interested if this promoter was in general functional in *C. glabrata* isolates. As such, we generated a Nourseothricin-based vector backbone (with the ScCEN/ARS origin of replication) containing the MET3p-Venus-ENOt transcriptional unit and used it to transform two different isolates of *C. glabrata*. Expression changed 4 and 5-fold (in each isolate respectively) when using methionine as repressor and showed no significant leakiness in the presence of >0.625 μM methionine (when compared to background**) (Supplementary Figure 11)**. Similar as observed in ATCC 2001 HTL^−^, cysteine was not as effective in repressing the promoter but allowed for a wider dynamic range than methionine.

### *S. cerevisiae* ubiquitin-based degradation tags can be ported into *C. glabrata*

We further tested if the ubiquitin-based YTK-derived degradation tags could be functionally ported into *C. glabrata*. In addition to controlling transcript levels via promoter strength, protein levels and turnover rates can be tuned by fusing degradation tags to the N-terminus of a protein.^44^ Perturbing turnover rates can be used for functional proteomics studies.^45^ The YTK-derived degradation tags UBI-M (weak), UBI-Y (medium) and UBI-R (strong) were fused to the Venus protein cloned under the control of the TDH3 and ALD6 promoters. As before, expression levels were measured in bulk and normalized to fluorescein. **Figure 2G** shows that all three degradation tags were functional in *C. glabrata* reducing the fluorescence in cells with a TDH3-driven degradation-tagged Venus to 56% (UBI-M), 39% (UBI-Y), and 37% (UBI-R) of the untagged Venus levels. For the ALD6p-driven degradation-tagged Venus, fluorescence was reduced to 52% (UBI-M), 33% (UBI-Y), and 20% (UBI-R). In comparison, in *S. cerevisiae* the tagging led to a remaining fluorescence of 63% (UBI-M), 30% (UBI-Y) and 15% (UBI-R) for a TDH3p-driven degradation-tagged Venus protein and to 92% (UBI-M), 36% (UBI-Y) and 23% (UBI-R) for an ALD6p-driven degradation-tagged Venus protein **(Supplementary Figure 12)**.

### A CRISPRi system for use in *C. glabrata*

Finally, we used the CgTK to build and test a CRISPRi system for *C. glabrata*. The system is based on the deactivated Cas9 (dCas9) fused to the transcriptional repression domain Mxi1,^46^ in combination with the gRNA design featured in the YTK. This gRNA design uses a gRNA fused to a self-cleaving hepatitis delta virus (HDV) ribozyme to enhance gRNA abundance (CRISPRm design).^47^ We first cloned a BsaI/BsmBI restriction site-free dCas9-Mxi1 construct as a level 0 vector and verified that this recoded fusion would be functional by testing it in *S. cerevisiae* with an established gRNA/promoter pair.^48^ We then cloned the dCas9-Mxi1 fusion protein under the control of an HHF1 promoter and ENO1 terminator on vector version 1.3 (LEU2, ScCEN/ARS). The gRNAs were expressed from a second vector, vector version 2.2 (TRP1, CgCEN/ARS). We chose to test repression of genes by a single gRNA as we are most interested in using the CRISPRi system for functional genomics studies based on single-gRNA library screens. Single-gRNA library screens are based on the fact that single gRNAs repress a target gene enough to generate selectable phenotypes such that cells carrying a given phenotype can be enriched or depleted from a library over various growth and dilution cycles.

We used two test genes to show the functionality of our CRISPRi system: (1) The repression of the *URA3* gene (CAGL0I03080g) and (2) the repression of the *ALG3* gene (CAGL0A04587g). The repression of the *URA3* gene should lead to a selectable phenotype on media lacking uracil. *ALG3* is a functionally yet unverified gene in *C. glabrata*, but its *S. cerevisiae* orthologue encodes for an alpha-1,3-mannosyltransferase, an endoplasmic reticulum localized enzyme involved in (membrane)-protein glycosylation. An *ALG3* deletion renders *S. cerevisiae* resistant to the yeast killer toxin HM-1. As we are interested in using the CRISPRi system to study resistant phenotypes to yeast killer toxins in *C. glabrata*, we verified that an *ALG3* deletion would also render *C. glabrata* resistant to this toxin **(Supplementary Figure 13)** and subsequently used it as a test target to verify that its repression by CRISPRi also changes the resistance profile.

We designed four to five gRNAs targeting the promoters of *URA3* and *ALG3* in a window of -50 to + 300 bp of the transcriptional start site (TSS) **(Supplementary Table 5)**. TSSs in *C. glabrata* have been determined experimentally^49^ and are available via the candida genome base.^50^ We chose this design as for *S. cerevisiae* it was shown that the most effective gRNAs are those designed to target a 50 to 200 bp window upstream of the TSS.^51,52^ For mammalian cells, effective gRNAs target a window of -50 to +300 bp^53^ around the TSS.

First, we tested the *URA3*p targeting CRISPRi system by measuring growth in media with and without uracil. For three out of the five gRNAs, we observed a slow growth phenotype in the absence of uracil **(Figure 4A and B)**, which allowed to deplete the gRNA-encoding strains over three growth-and-dilution cycles to 4 to 10-fold (depending on gRNA) when compared to the same strain grown in the presence of uracil or a strain not expressing a targeting gRNA **(Figure 4C)**. Next, we assessed the performance of the *ALG3*p targeting CRISPRi system: Here one out of the four tested gRNAs could be enriched in the presence of the killer toxin HM-1 **(Figure 4D and E)**.

**Figure 4.**
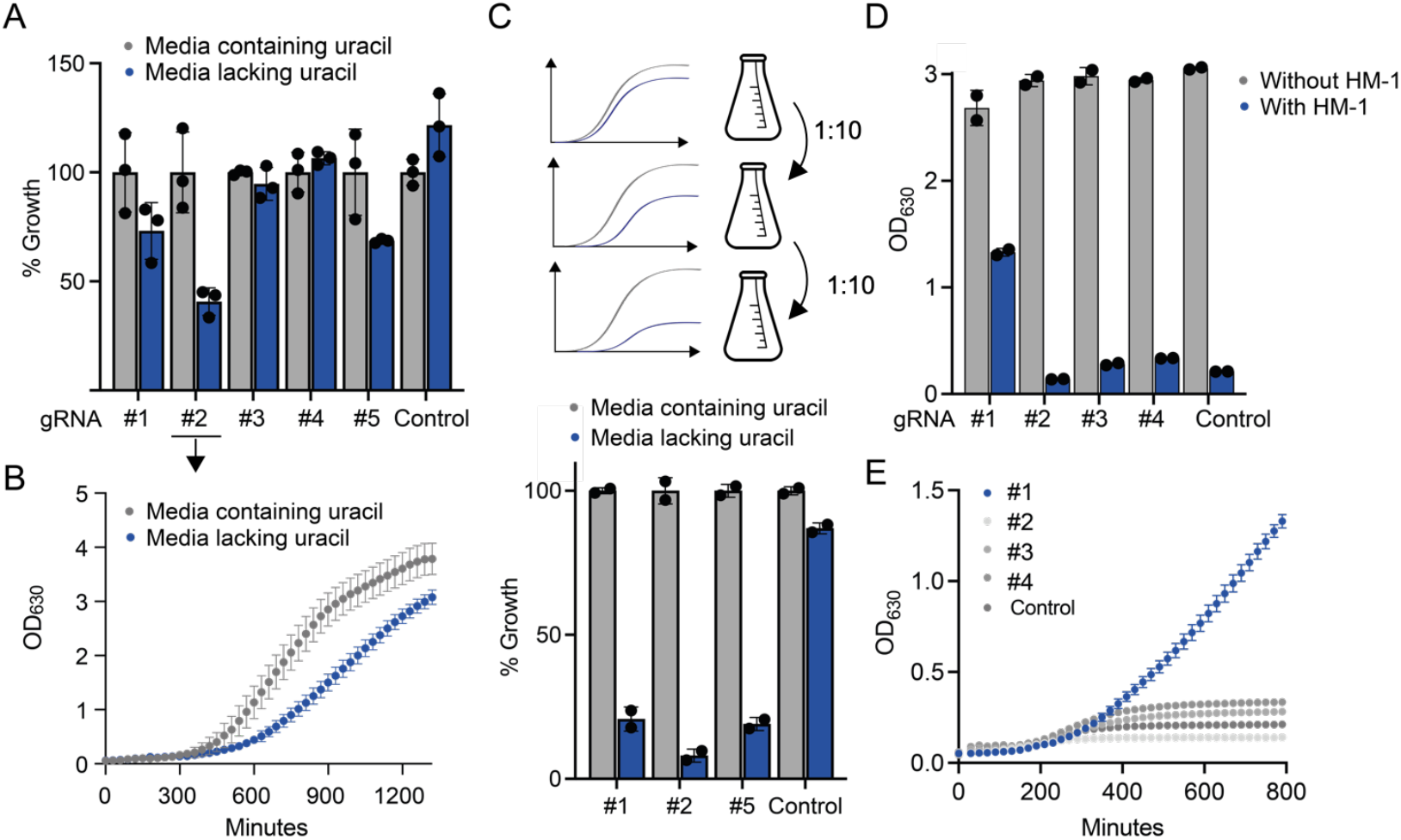
CRISPRi in *C. glabrata*. **A:** Two test systems were used, single-gRNA repression of *URA3* **(A-C)** and *ALG3* **(D-E). A:** Five gRNAs were tested for repression of the *URA3* gene by measuring growth in media lacking uracil. Percent growth (OD_630_) after 12 hours when compared to the same strain grown in media with uracil is depicted. The control uses a gRNA that does not target the *URA3* promoter. **B:** Full growth curve of the strain harboring the best performing gRNA #2 in media containing uracil and media lacking uracil. **C:** Depletion of strains harboring gRNAs #1, #2, and #5 after three serial dilutions in media lacking uracil when compared to growth in media containing uracil. Strains were grown in the presence or absence of uracil in duplicate and cultures were diluted 1:10 into fresh media after seven to ten hours. Final OD_630_ after the third dilution round is given as percentage of growth in media containing uracil. **D:** Four gRNAs were tested for repression of the *ALG3* gene by measuring growth in media containing the killer toxin HM-1. The control uses a gRNA that does not target the A*LG3* promoter. **E:** Growth curves of the strains harboring the targeting or non-targeting RNAs in media containing HM-1.

## CONCLUSION

Here we deliver a molecular cloning kit for the human fungal pathogen and biotechnology host *Candida glabrata* by characterizing the performance of the YTK and by adding several new parts. In total, the kit contains 25 constitutive promoters (19 YTK, 2 from *C. glabrata* and 4 minimal promoters), 3 inducible/repressible promoters (copper and methionine/cysteine), 3 degradation tags, and 9 pre-assembled vectors with different selection markers.

The strongest promoters generated expression levels 100-fold over background and allow a user for tuning expression levels over two orders of magnitude. The herein characterized vector-set allows for further tuning expression strength as they have different copy number.

To further enhance expression strength of the promoters, it might be possible to choose better suited “seams” for the Golden Gate assembly of type 2 (promoter) and type 3 (open reading frame) parts as the YTK-inherent BglII site was shown to be suboptimal.^43^

While here we show a single-gRNA CRISPRi system useful for library selections, the system could likely be extended to generate complete knock-down phenotypes (as required for metabolic engineering) by using several (3 or more) gRNAs that target the same promoter and that need to be expressed in the same cell. YTK compatible multi-gRNA cloning strategies are readily available to extend the herein presented single-gRNA system.^48^

In summary, we think that the CgTK is a starting point for effective metabolic engineering and phenotypic characterization of *C. glabrata*.

## MATERIALS AND METHODS

### Materials

Phire Green Hot Start II PCR Master Mix was used for all PCR reactions and was purchased from Thermo Fisher (#F126L). Restriction enzymes BsaI-HF®v2 (#R3733S) and BsmBI-v2 (#R0739S) and T7 ligase (#M0318S) were used for the Golden Gate reactions and were obtained from New England Biolabs (NEB). Media components were obtained from BD Bioscience and Sigma-Aldrich. Primers and synthetic DNA (gBlocks) were obtained from Integrated DNA Technologies (IDT); Primers used in this study are listed in **Supplementary Table 4**. Plasmids were cloned and amplified in *E. coli* DH5α. Clear, round bottom 96-well microtiter plates (Costar) were used for culturing. Black, clear-bottom 96-well microtiter plates (Costar) were used for fluorescence measurements. Fluorescence and optical density measurements were performed in a SynergyMx (Biotek) plate reader at 630 nm (optical density), 561ex/610em (red fluorescence) and 488ex/530em (green fluorescence).

### Strains

Experiments were either performed in *C. glabrata* ATCC 2001 HTL or its derivative HTLU^−^ and in *S. cerevisiae* BY4741 (*MATa leu2Δ0 met15Δ0 ura3Δ0 his3Δ1*).^54^ *E. coli* DH5α was used for cloning. Strain *C. glabrata* ATCC 2001 HTL^−^ DeltaRHK1/ALG3^38^ (ORF CAGL0A04587g deleted) was used as a control for the ALG3 CRISPRi experiments.

### Plasmids, primers and synthetic DNA

Plasmids are listed in **Supplementary Table 1 and 2**, sequences of primers are listed in **Supplementary Table 4**, synthetic DNA (gBlocks, IDT) are listed in **Supplementary Table 3**.

### Media and inductions

Yeast strains were routinely grown in synthetic drop-out media (SD) with 2% (w/v) Dextrose (Fisher Scientific), 0.67% (w/v) Yeast Nitrogen Base without amino acids (VWR International), and a self-made 20-fold amino acid drop-out mix.^55^ Nourseothricin was used at 100 μg/mL final concentration in liquid or solid YPD media (yeast extract 2%, peptone 1%, dextrose 2%).

*E. coli* was grown in Luria Broth (LB) media. To select for *E. coli* plasmids with drug-resistant genes, ampicillin (Sigma-Aldrich) or kanamycin (Sigma-Aldrich) were used at final concentrations of 75-200 μg/mL and 50 μg/mL, respectively. Agar was added to 2% for preparing solid yeast and bacterial media. Copper inductions were performed in SD media with 0-2 mM copper (II) sulfate added. Methionine/cysteine repressions were performed in SD media lacking methionine in the drop-out mix, with 0-10 mM methionine or cysteine added.

### Yeast transformation

*C. glabrata* and *S. cerevisiae* cells were transformed with plasmid DNA using the lithium acetate transformation protocol described before.^56^

### Cloning of new parts into the entry vectors (pYTK01) and Golden Gate assembly

Primers with overhangs for BsmBI-based cloning into the entry level vector pYTK01 were designed as instructed in Lee *at al*.^29^ Golden Gate reactions were performed as instructed in Lee *at al*.^29^ In brief: 0.5 μL of each DNA insert or plasmid, 1 μL T4 DNA Ligase buffer (NEB), 0.5 μL T7 DNA Ligase (NEB), 0.5 μL restriction enzyme, and water to bring the final volume to 10 μL. The restriction enzymes used were either BsaI or BsmBI (both 10 000 U/mL from NEB).

Reaction mixtures were incubated in a thermocycler according to the following program: 25 cycles of digestion and ligation (42 °C for 2 min, 16 °C for 5 min) followed by a final digestion step (60 °C for 10 min), and a heat inactivation step (80 °C for 10 min). In some cases, where noted in the text, the final digestion and heat inactivation steps were omitted.

On a technical note; Golden Gate-based cloning of dCas9-MxiI was less reliable and at least 5 white colonies needed to be picked after green white screening and their plasmids control digested in order to identify one correct assembly. This is in contrast to the usually highly efficient YTK-based Golden Gate assembly were almost 100% of white colonies carry correct assemblies.

### Fluorescence measurements and fluorescence calibration

Three colonies were picked and grown in 200 μL of media in 96-well plates (clear, round bottom) at 30°C in an orbital shaker (microtitre plate shaker incubator SI505 from Stuart) shaking at 950 rpm until saturated. Cultures were diluted 1:100 in fresh media, grown for 16 h at the same conditions as the overnight cultures, and then diluted 1:2 in water in 96-well black clear-bottom plates, and fluorescence was measured in a SynergyMx plate reader. For the copper inductions, saturated cultures were diluted 1:100 in fresh media with different concentrations of copper (II) sulfate and grown for 16 h. For the methionine/cysteine repression, saturated cultures (grown in the presence of methionine, OFF state) were washed 5 times in 5 volumes of water and diluted 1:100 in fresh media with different concentrations of methionine or cysteine added and grown for 16 h.

Excitation and emission wavelengths used to measure fluorescent proteins were 588 nm/530 nm for Venus and 561 nm/610 nm for mRuby2. Raw fluorescence values were first corrected by subtracting the average autofluorescence of a black clear-bottom plate, then normalized to the OD_630_ of the cultures, and then normalized to the respective calibrant dye (fluorescein for Venus and sulforhodamine-101 for mRuby2). Calibrating arbitrary fluorescence units into nM calibrant dye fluorescence was performed as described before.^39^ Linear regression was used to derive the conversion factor.

### Growth measurements

Growth curves were recorded in sterile, transparent round-bottom 96-well plates using 200 μL total culture volume, cultured at 30°C in a SynergyMx plate reader (high orbital shaking). Cells were seeded at an OD_630_ of approximately 0.03 and culture turbidity (OD_630_) was recorded every 30 minutes for 20 to 24 h. For gRNA depletion experiments, cells were grown in 50 mL shake flasks and since the later optical density values were outside the linear range of the photodetector, all optical density values were first corrected using the following formula to calculate true optical density values:

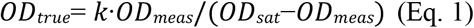

where *OD*_*meas*_ is the measured optical density, *OD*_*sat*_ is the saturation value of the photodetector (2.568 for our instrument, as experimentally determined) and *k* is the true optical density at which the detector reaches half saturation of the measured optical density (2.075 for our instrument, as experimentally determined).

For gRNA depletion experiments shown in **Figure 4C**, cells were grown in 50 mL shake flasks and diluted 1:10 every 7 to 10 h. Optical density was measured in 1 mL cuvettes in a spectrophotometer (Nanospec, Amersham Bioscience). For selections in the presence of HM-1, HM-1 was concentrated from an in-house production strain and used as 0.4x diluted supernatant in the final assay.

### Plasmid/DNA extraction and qPCR

Two different protocols for extracting DNA or plasmids from *C. glabrata* were employed:

#### Plasmid extraction

4 mL of media were inoculated with a *C. glabrata* colony harboring the targeted plasmids and incubated at 30 °C shaking at 200 rpm for 16 – 18 h. The equivalent of 5 OD units were used for extraction. The cells were centrifuged at 10,000 g for 15 sec and resuspended in 1 M Sorbitol, 100 mM EDTA, 14 mM β-mercaptoethanol, 20 U/mL lyticase (Sigma #L2524-10KU) for 1 h. For further processing the GeneJET Plasmid Miniprep Kit (Thermo Fisher) was utilized as instructed. Plasmid DNA was eluded in 30 μL of water.

#### Total DNA extraction from C. glabrata

For total DNA extraction 4 mL of media were inoculated with a *C. glabrata* colony harboring the targeted plasmids and incubated at 30 °C shaking at 200 rpm for 16 – 18 h. The equivalent of 10 OD Units used for DNA extraction. For this the cells were pelleted and washed in 50 mM EDTA and subsequently incubated in 50 mM EDTA, 20 U/mL lyticase (Sigma #L2524-10KU) at 30°C for 1 h. The pellet was resuspended in 50 mM EDTA, 2% SDS and incubated at 65 °C for 15 min before adding 1:1 potassium acetate. After centrifugation at 20,000 *g* for 10 minutes the supernatant was transferred to a fresh tube and isopropanol was added 1:1. The solution was again centrifuged at 20,000 *g* for 10 min, after which the DNA pellet was washed with 70% EtOH, air-dried and resuspended in 25 μL of water.

#### qPCR

qPCR reactions containing 4 μL sample, 12.5 μL 2x SensiFAST SYBR Hi-ROX Mix (Meridian Bioscience), 1 μL forward and reverse Primer (10 mM**) (**see **Supplementary table 4** for primer sequences) and water for a total volume of 25 μL was performed in white 96-well plates. Relative copy numbers and relative normalized copy numbers were determined in three biological replicates, which were measured in three technical replicates respectively. Standard cycling conditions as recommended by the manufacturer of the reaction mix were employed for data generation. For determining relative normalized expression of plasmid per cell, total DNA extract sample was added to the reaction mixture. The amount of measured plasmid (primers Venus-fw and Venus-rev) was normalized for the amount of actin (Actin-fw and Actin-rv) in each sample and the average of each technical replicate group was calculated and again averaged within the biological group resulting in an average ΔCq value per biological replicate group. The ΔCq values of each biological replicate group were then cross-compared yielding ΔΔCq values. The fold-changes corresponding to these ΔΔCq values are shown in **Supplementary Figure 4B**. For relative copy numbers samples of plasmid extraction were added to the reaction mixture. The amount of plasmid per sample was measured (primers Venus-fw and Venus-rev) and the relative amount determined without normalizing for actin by directly averaging the technical and thereafter the biological replicates. The fold-changes corresponding to the ΔCq values of the cross-comparison between the biological replicate groups **Supplementary Figure 4A**.

## Author contribution

SB designed the work, performed experiments and wrote the paper. RCP cloned and tested the BsaI- and BsmBI-site free dCas9-MxiI. MM performed qPCR experiments.

## Acknowledgement

We thank Prof. Karl Kuchler (Medical University Vienna) for providing strains ATCC2001 HTL and ATCC 2001 HTL^−^ DeltaRHK1

