## Supplementary Material for "A modular cloning toolbox including CRISPRi for the engineering of the human fungal pathogen and biotechnology host *Candida glabrata*"

Sonja Billerbeck *et al.*

Molecular Microbiology, Groningen Biomolecular Sciences and Biotechnology Institute, University of Groningen, Groningen, The Netherlands

**Supplementary Table 1.** YTK parts that were used for vector assembly or re-characterized in *C. glabrata.*

| YTK# | Name | Type |
|---|---|---|
| YTK parts re-used for vector backbones | | |
| pYTK08 | ConLS' | 1 |
| pYTK73 | ConRE' | 5 |
| pYTK74 | URA3 | 6 |
| pYTK75 | LEU2 | 6 |
| pYTK76 | HIS3 | 6 |
| pYTK78 | NourseothricinR | 6 |
| pYTK84 | KanR-ColE1 | 8 |
| pYTK81 | CEN6/ARS4 | 7 |
| pYTK47 | GFP dropout | 2/3/4 |
| YTK promoters characterized in *C. glabrata* | | |
| pYTK09 | pTDH3 | 2 |
| pYTK10 | pCCW12 | 2 |
| pYTK11 | pPGK1 | 2 |
| pYTK12 | pHHF2 | 2 |
| pYTK13 | pTEF1 | 2 |
| pYTK14 | pTEF2 | 2 |
| pYTK15 | pHHF1 | 2 |
| pYTK16 | pHTB2 | 2 |
| pYTK17 | pRPL18B | 2 |
| pYTK18 | pALD6 | 2 |
| pYTK19 | pPAB1 | 2 |
| pYTK20 | pRET2 | 2 |
| pYTK21 | pRNR1 | 2 |
| pYTK22 | pSAC6 | 2 |
| pYTK23 | pRNR2 | 2 |
| pYTK24 | pPOP6 | 2 |
| pYTK25 | pRAD27 | 2 |
| pYTK26 | pPSP2 | 2 |
| pYTK27 | pREV1 | 2 |
| pYTK31 | pCUP1 | 2 |
| YTK fluorescent proteins characterized in *C. glabrata* | | |
| pYTK33 | Venus | 3 |
| pYTK34 | mRuby2 | 3 |
| YTK terminators characterized in *C. glabrata* | | |
| pYTK51 | tENO1 | 4 |
| YTK degrons characterized in *C. glabrata* | | |
| pYTK41 | Ubi-M | 3a |
| pYTK42 | Ubi-Y | 3a |
| pYTK43 | Ubi-R | 3a |

**Supplementary Table 2.** New parts added to the *Cg*TK.

| **CgTK#** | **Name** | **Part type** | **Source of DNA and reference** |
|---|---|---|---|
| **Level 0 parts** | | | |
| CgTK parts for creating vector backbones | | | |
| pCgTK01 | *Cg*CEN/ARS | 7 | pCU-Met3-GFP (Addgene# 45337)[1] |
| pCgTK02 | CgTRP1 | 6 | Genomic DNA of ATCC2001, BsaI/BsmBI sites removed |
| CgTK promoter parts | | | |
| pCgTK03 | CgMET3p | 2 | pCU-Met3-GFP (Addgene# 45337)[1] |
| pCgTK04 | CgMT1p | 2 | Genomic DNA of ATCC2001 |
| pCgTK05 | CgTDH3p | 2 | Genomic DNA of ATCC2001 |
| pCgTK06 | CgPGK1p | 2 | Genomic DNA of ATCC2001 |
| pCgTK07 | MP1 (Core 1, UAS c-e-f) | 2 | Ordered as synthetic DNA, design derived from Redden *et al.*[2] |
| pCgTK08 | MP2 (Core 5, UAS c-e-f) | 2 | |
| pCgTK09 | MP3 (Core 8, UAS c-e-f) | 2 | |
| pCgTK10 | MP4 (Core 9, UAS c-e-f) | 2 | |
| CgTK *CRISPRi* parts | | | |
| pCgTK11 | NLS-dCas9-Mxi1 (BsaI- and BsmBI-site free) | 3 | Ordered as synthetic DNA, design based on Smith *at al.*[3] The original construct contained 3 BsaI sites and 1 BsmBI site. We modified these 4 regions to remove all BsaI and BsmBI sites. |
| **Preassembled vectors** | | | |
| CgTK vectors for cloning (multiple) TUs | | | |
| CgTK13 | Version 1.1 (ScCEN6/ARS4, *HIS3*) | | |
| CgTK14 | Version 1.2 (ScCEN6/ARS4, *TRP1*) | | |
| CgTK15 | Version 1.3 (ScCEN6/ARS4, *LEU2*) | | |
| CgTK16 | Version 1.4 (ScCEN6/ARS4, *URA3*) | | |
| CgTK17 | Version 1.5 (ScCEN6/ARS4, *NAT1*) | | |
| CgTK18 | Version 2.1 (CgCEN/ARS, *HIS3*) | | |
| CgTK19 | Version 2.2 (CgCEN/ARS, *TRP1*) | | |
| CgTK20 | Version 2.3 (CgCEN/ARS, *LEU2*) | | |
| CgTK21 | Version 2.4 (CgCEN/ARS, *URA3*) | | |
| CRISPRi vectors | | | |
| CgTK22 | Vector 2.2_HHF1p_dCas9-Mxi1_ENO1t | | |

**Supplementary Table 3.** Sequences of new *Cg*TK parts. The part-specific overhangs are indicated in bold, green. The sequences for the minimal promoter units are annotated as follows: Capitals, green: upstream activating sequence c; Capitals, dark blue: upstream activating sequence e; Capitals, red: upstream activating sequence f; Bold, black: minimal core promoter; Blue: spacer.

| CgTK number | Description | Sequence |
|---|---|---|
| pCgTK01 | *Cg*CEN/ARS | **GAGT**ttgcagtcgtactggatctgtgaatctattagtatatatgaattaaagtagcttgtacatattatt<br>ctgttgaatcatatcgcagagcattggttgaaatccaaaatataaaaatgtaatatcacaaaaaataata<br>ctaattctaacattaatgggtcagatttttagtgaatacttaaatttataatatcgtctatttaagctagcaaa<br>tggaacaacatttaaagtaagaacatcatatctacatgaaaatgtatatttcaatctgactaataacgca<br>gagcacatctttcagtgatgtctgtcacatgatcaaaaagaattgtatttaaataattcataataaaagctt<br>aaaaaattacaataatgaaaataaagtaaataatgacatgggtaagagttccgaaatagaatcttagtg<br>tacaaaacaaaaattgcatcattagagatcccca **CCGA** |
| pCgTK02 | Cg*TRP1*p-<br>*TRP1*-*TRP1*t | **TACA**agtgcaaaggcatttcctttgccacacacactaccataactagtctcggtattaactggctca<br>acaaacgataagagccaaagaggaagaaaaaaaaggtatttaaaggatgtatttctttatctcctgatt<br>agatttcttacttgtacagaaactgtaactaaaaaaaaacaacaacaaaacacaacacatacataatgg<br>ctagaaccaagcaaaccgcaagaaagtctactggtggtaaggccccaagaaagcaactagcttcta<br>aggctgccagaaaatccgctccatctaccggtggtgtcaagaagcctcacagatataagccaggtac<br>cgtcgctttgagagaaatcagaagattccaaaagtctactgaacttttgatcagaaagttgcctttccaa<br>agactagtcagagaaatcgcccaagatttcaagaccgatctaagattccaatcctctgccatcggtgc<br>tctacaggaatccgttgaagcttacttagtctctttgttcgaagacaccaacttggccgctatccacgct<br>aagcgtgttaccatccaaaagaaggatatcaagttggctagaagattgagaggtgaaagatcctaag<br>cacacagggattgttgtacaattactttttatgattttagatacacacaaatcttttctacgattacctttcatt<br>tattttcgtttcatgttttattaatcattctttttgtatatttcacatcgggtggggtttttatattaaattttaataa<br>cataatcttttaaagtttctactttctatcctgtatattatttccatatctatatctatattttcaagcttgttcaca<br>tccttatcggaattgttccagcgtttttgtacagaaaatgatgttatggtaacgagctatatcagacctact<br>agtcttatagagctgtacagagtcttggaaacataaccaacacaccaacaagatgtcatttgattcgtta<br>ctcgacaagaatgataagctggtaaaagtttgcgggattcaaaccgtcgaggctgccgaaactgcgc<br>ttcaagcaggcgctgatttgatagggatcatatgtgtccccaacaggaagcggactatcgagagcgc<br>tgtggctcgtgaaatatccaaattgattcacaaatcaggtactacaaagctggtgggggtgttcagga<br>atcaatctgttgaggacgtacatcggctttctgaggaatatgaccttgacataatccaattacatggtga<br>tgaatcatggccagagtactataacgtcattaagaaaccaataatcaaaagagtcatattccctagaga<br>tgtcgatgttgtaacacaagtgtgtcaaagaaaacccttggtatgtctaccattgttcgactccgaggca<br>ggcggtacaggtgaaaagctggactggtccagtatttcctcctgggcaagcgaacagaataacgtat<br>tttatatactagcgggtggactaactgcagaaaacgtcatggaggcggtggggttgcctggtgtaatt<br>ggtgttgacgtatcaggtggtgtagagactgacggcgtcaaagacaatgacaagataattaagtatgt<br>acaaaatgcaaagaaacaatgaattacgaagagaagtctcaatgaggttttacaaacaatcctataat<br>acatatatgccactatgatatactttgaatactgagtatactgactcattatattctttaattgtagtttctcaa<br>aaatattcatacaggctagctatgaattcatgctataatttccattttgttcatgtaagatacacatagggga<br>taaattgtgataactaatcagccgacttccgggtaccgttaatattatgcgtacttgttttcatgcgttgcg<br>ttattcaccctctttttaaatcgtcagtcgaattcgtaaacttcttaaataatatcgatattgattacatctgttt<br>agtaatcaaaacattcgataagtcagaattcattcaatgcataaaaatctgtgaatactgaggaacaatt<br>ggagtttcagtaaaaagtcttgcaaagcatgccaagttcagtgactgtcttacacatattaccacagaa<br>gtatcttcaccaaatttagtc**GAGT** |
| pCgTK03 | CgMET3p | **AACG**ggcttatatgtatggaggtagggaactgcagtttattgttctgttaagctcccagtaatgcaa<br>gacggctaaatcatatgactgccactttgttgatcctgaagaaaatgacaacaagtagaaagtgatata<br>atacacgtggtatccgtagtatggtatcaggagatagccacatcacatgatctctaaaaccccccgtc<br>agcaaggaataaatcgaagagaaaaaaatgccacgtgactttgatggctaaaatcaggttatactact<br>gtacggtcccccacacataacctttaccacacaccgcacgggctggactcatcctattagaaagcgg<br>tgcagccagggccaagaaaacgcgaacgacgcagaaaaaacgtcaagcaaaaaaactgtggtgtt<br>tttttaaagcatacaatttcctgcctccctttaatgctacggggatctagcaaatgggaaaatatcacatg<br>actttggctagaaggggtgggaaaacagagattttagtcacattgtttgatttcacgtactacacgatac |

| | | |
|---|---|---|
| | | actacactacacaatacactccagtgcaatacactccagtgcaatacactccagtgcaatacactccagtgcaatacactccagtgcaatacactgcaatacactacactgtatggttccctccccgtcctcttcaggccctcgatatgctagcgaaggatcccaagcccatcgaggaaatcattcaagggcatctgcaggtctcaaatacactaagtccaaacacaaacagcactgagctaatccaagacaccaattgcatgccttctcaattatatcagtttattacacagcttataactgtgcatcttgccattcttttcgagatagccatttgattaacatgcttagctcgttctcaagccacagtaaaatggattgccttttgagtttccatcgttgtataaataggctctcattcctatagccttgctctttggtcttgtctcggtaatagtaatagctgtaagtcaggatataacactccagaaaagaaacacctaacaa**TATG** |
| pCgTK04 | CgMT1p | **AACG**tctggaacttctgggaactcgctcccacacagaagaaatcgcaccgtgctagaccggatcagaagggatatcagcagggatcgcagtcgcggagtaacggccgtacacactctctcgtcacaactggaaagaacaaccgacgggccggatcggaacataaattgctttttcctgcccttcctttttctgatgctcatacgtgctgctctgtgcttgcagcgcaatttgtagatctgttgtacgcgtgtttctcaagcaccaacccagtcaactttgagcttaactgcctcactattaataatatccgcatctcacaaataaccaattttgctgattctgtctagtgccacttgtcacttgccagtgccagtgccagtgctcacttgccaggtttgctgcttaaagaattccgctcttcatacacatcctacactcccttaaggagggaagtacaaacaagaagtgttcgcccattctctctcacatgatgtgtaaagaaaagtgaaaatcagcaataatagcttctgactatgactatctgtattgacagcaataatagctttcgggatagcatgcaaaaaaaaatttatatataaacagaggtctttttgaaatgttagatcaatttctaattaaacattttgtagatcctacataactcatacaacaaacaacaaacacatacaaaaacaaacaca**TATG** |
| pCgTK05 | CgTDH3p | **AACG**tcattcaacaaacaccttttctttagatcttgcttttgttattacccttactattgtctggctctgcgccattctttccactaacaatacgggtgcatggcaatcattttatgaaaatatatacacatacatctcatgataaatttcagttgtagcctcccatcaagagttagttaaaccacatgtatagctgtctctaggaataatgaaatgataatgcaggcccgagactgtattattgtgtctttggcaccttaaaaaaaagtgtcttccctccatagtgtgcacctacaaaaaatttacatacatatcccatagtgctaacaacttccttagataagacaaaactctatagtgaccaagaacagtattatcaaaattcaataacctgcaatatgacaattcccaacaaagagacaattatcaattctagaataactcttgtgcattttacactcaactgaagaggtattaatccctctcaagaagcaaacagcataataggatatgcctcccagcttcaaaataatcatataaatatgtgccctctttgcaagtaactgacatcaattgtccttgaattcattccattttaaacattatctataaagcttgaaaaaaccacagctacaagtattaattactacaggctaaagatacacagaacacatacacataacaaaacttattaacttcaaaa**TATG** |
| pCgTK06 | CgPGK1p | **AACG**acacggctgtggtcgtcgtgacgaccacagccgtgtataataacaaaaagcagttaacaaataatagcaaacggtacacatgtgactaccacgtattgtatgctttacgggtgacaagaaaaggaaaccccgccaagatacgtgcaagacgagaataccagctttctgcttttctttcttcttcctcgatttaatggctgggggggaactgtgagattaccatgacacccgtgcacccccggactccactaggggaagcatccaggtcgggtgggagtgagagggggaatcgggcaactgctctaggagaagctctaggagaagctctaggagaagccctaggagaagcaaaggggaagcccaaagagggacaagcgcaggccactacagtaacaaagacaggaagcccaaaagatcgcaccttccagtccatcgccttcccccattgtcatatggcagtttatagcagtgtggcagtatggcagtgtggcaatgtgaagcagtgtgcaatacttccaaagtggataatgtgatgcttgcaaaaagttgggaccaccaatgctgtagagacaatgggataccctagcgaatgaagaaaaaaaggaaaacgaaaaaaatttgtatataaagcagaggcaatttttgtagcggcatgattagttgtcaattggcaagacggcatacatacatctattcgata**TATG** |
| pCgTK07 | Core 1 UAS c-e-f | ggcgcgccCCTCCTTGGAACTGAAATTTTAGCATGTGAttaattaacttgtaatattctaatcaagct**tataaaagagcactgttgggcgtgagtggaggcgccgg**aaaaaagcatcgaaaaaatctag |
| pCgTK08 | Core 5 UAS c-e-f | ggcgcgccCCTCCTTGGAACTGAAATTTTAGCATGTGAttaattaacttgtaatattctaatcaagct**tataaaagggccttggtctgaaactcctgcgtctcgcg**aaaaaagcatcgaaaaaatctag |
| pCgTK09 | Core 8 UAS c-e-f | ggcgcgccCCTCCTTGGAACTGAAATTTTAGCATGTGAttaattaacttgtaatattctaatcaagct**tataaaagcaatacttgggtcgacttgttatacgcgga**aaaaaagcatcgaaaaaatctag |
| pCgTK10 | Core 9 UAS c-e-f | ggcgcgccCCTCCTTGGAACTGAAATTTTAGCATGTGAttaattaacttgtaatattctaatcaagct**tataaaagggcgctgcgtaaggagtgctgccaggtggt**aaaaaagcatcgaaaaaatctag |

| | | |
|---|---|---|
| pCgTK11 | NLS-dCas9-<br>Mxi1 | **TATG**tctagagccccaaagaagaagagaaaagttagacccgggatggacaagaagtactccat<br>tgggctcgctatcggcacaaacagcgtcggctgggccgtcattacggacgagtacaaggtgccga<br>gcaaaaaattcaaagttctgggcaataccgatcgccacagcataaagaagaacctcattggcgccct<br>cctgttcgactccggggaaacggccgaagccacgcggctcaaaagaacagcacggcgcagatat<br>acccgcagaaagaatcggatctgctacctgcaggagatctttagtaatgagatggctaaggtggatg<br>actctttcttccataggctggaggagtcctttttggtggaggaggataaaaagcacgagcgccaccca<br>atctttggcaatatcgtggacgaggtggcgtaccatgaaaagtacccaaccatatatcatctgaggaa<br>gaagcttgtagacagtactgataaggctgacttgcggttgatctatctcgcgctggcgcatatgatcaa<br>atttcggggacacttcctcatcgagggggacctgaacccagacaacagcgatgtcgacaaactcttt<br>atccaactggttcagacttacaatcagcttttcgaagagaacccgatcaacgcatccggagttgacgc<br>caaagcaatcctgagcgctaggctgtccaaatcccggcggctcgaaaacctcatcgcacagctccc<br>tggggagaagaagaacggcctgtttggtaatcttatcgccctgtcactcgggctgacccccaacttta<br>aatctaacttcgacctggccgaagatgccaagcttcaactgagcaaagacacctacgatgatgatctc<br>gacaatctgctggcccagatcggcgaccagtacgcagaccttttttggcggcaaagaacctgtcag<br>acgccattctgctgagtgatattctgcgagtgaacacggagatcaccaaagctccgctgagcgctag<br>tatgatcaagcgctatgatgagcaccaccaagacttgactttgctgaaggcccttgtcagacagcaac<br>tgcctgagaagtacaaggaaattttcttcgatcagtctaaaaatggctacgccggatacattgacggc<br>ggagcaagccaggaggaattttacaaatttattaagcccatcttggaaaaaatggacggcaccgagg<br>agctgctggtaaagcttaacagagaagatctgttgcgcaaacagcgcactttcgacaatggaagcat<br>cccccaccagattcacctgggcgaactgcacgctatcctcaggcggcaagaggatttctacccctttt<br>tgaaagataacagggaaaagattgagaaaatcctcacatttcggataccctactatgtaggcccctc<br>gcccggggaaattccagattcgcgtggatgactcgcaaatcagaagaaaccatcactccctggaact<br>tcgaggaagtcgtggataagggggcctctgcccagtccttcatcgaaaggatgactaactttgataaa<br>aatctgcctaacgaaaaggtgcttcctaaacactctctgctgtacgagtacttcacagtttataacgagc<br>tcaccaaggtcaaatacgtcacagaagggatgagaaagccagcattcctgtctggagagcagaaga<br>aagctatcgtggacctcctcttcaagacgaaccggaaagttaccgtgaaacagctcaaagaagacta<br>tttcaaaaagattgaatgtttcgactctgttgaaatcagcggagtggaggatcgcttcaacgcatccct<br>gggaacgtatcacgatctcctgaaaatcattaaagacaaggacttcctggacaatgaggagaacga<br>ggacattcttgaggacattgtcctcacccttacgttgtttgaagatagggagatgattgaagaacgcttg<br>aaaacttacgctcatctcttcgacgacaaagtcatgaaacagctcaagaggcgccgatatacaggat<br>gggggcggctgtcaagaaaactgatcaatgggatccgagacaagcagagtggaaagacaatcctg<br>gattttcttaagtccgatggatttgccaaccggaacttcatgcagttgatccatgatgactctctcacttt<br>aaggaggacatccagaaagcacaagtttctggccaggggacagtcttcacgagcacatcgctaat<br>cttgcaggtagcccagctatcaaaaagggaatactgcagaccgttaaggtcgtggatgaactcgtca<br>aagtaatgggaaggcataagcccgagaatatcgttatcgagatggcccgagagaaccaaactaccc<br>agaagggacagaagaacagtagggaaaggatgaagaggattgaagagggtataaaagaactggg<br>gtcccaaatccttaaggaacacccagttgaaaacacccagcttcagaatgagaagctctacctgtact<br>acctgcagaacggcagggacatgtacgtggatcaggaactggacatcaatcggctctccgactacg<br>acgtggatgctatcgtgccccagtcttttctcaaagatgattctattgataataaagtgttgacaagatcc<br>gataaaaatagagggaagagtgataacgtcccctcagaagaagttgtcaagaaaatgaaaaattatt<br>ggcggcagctgctgaacgccaaactgatcacacaacggaagttcgataatctgactaaggctgaac<br>gaggtggcctgtctgagttggataaagccggcttcatcaaaaggcagcttgttgagacacgccagat<br>caccaagcacgtggcccaaattctcgattcacgcatgaacaccaagtacgatgaaaatgacaaactg<br>attcgagaggtgaaagttattactctgaagtctaagctggtttcagatttcagaaaggactttcagttttat<br>aaggtgagagagatcaacaattaccaccatgcgcatgatgcctacctgaatgcagtggtaggcactg<br>cacttatcaaaaaatatcccaagcttgaatctgaatttgtttacggagactataaagtgtacgatgttagg<br>aaaatgatcgcaaagtctgagcaggaaataggcaaggccaccgctaagtacttcttttacagcaatat<br>tatgaatttttcaagaccgagattacactggccaatggagagattcggaagcgaccacttatcgaaac<br>aaacggagaaacaggagaaatcgtgtgggacaagggtagggatttcgcgacagtccggaaggtc<br>ctgtccatgccgcaggtgaacatcgttaaaaagaccgaagtacagaccggaggcttctccaaggaa<br>agtatcctcccgaaaaggaacagcgacaagctgatcgcacgcaaaaaagattgggaccccaagaa<br>atacggcggattcgattctcctacagtcgcttacagtgtactggttgtggccaaagtggagaaaggga<br>agtctaaaaaactcaaaagcgtcaaggaactgctgggcatcacaatcatggagcgatcaagcttcga<br>aaaaaaccccatcgactttctcgaggcgaaaggatataaagaggtcaaaaaagacctcatcattaag<br>cttcccaagtactctctctttgagcttgaaaacggccggaaacgaatgctcgctagtgcgggcgagct |

| | | |
|---|---|---|
| | | gcagaaaggtaacgagctggcactgccctctaaatacgttaatttcttgtatctggccagccactatgaaaagctcaaaggttctcccgaagataatgagcagaagcagctgttcgtggaacaacacaaacactaccttgatgagatcatcgagcaaataagcgaattctccaaaagagtgatcctcgccgacgctaacctcgataaggtgctttctgcttacaataagcacagggataagcccatcagggagcaggcagaaaacattatccacttgtttactctgaccaacttgggcgcgcctgcagccttcaagtacttcgacaccaccatagacagaaagcggtacacctctacaaaggaggtcctggacgccacactgattcatcagtcaattacggggctctatgaaacaagaatcgacctctctcagctcggtggagacagcagggctgacgagggagctcccaagaaaaagcgcaaggtaggtagttccaagcttggcggcagcggcggcagcatggaacgtgtgagaatgattaatgtgcaaaggctgttagaagccgcagagtttttagaaagaagagaaagagaatgcgaacacgggtatgccagttctttccctagcatgccctctcccagaggctaa**ATCC** |

**Supplementary Table 4**. Primers used in this study.

| Name | Sequence | Comment |
|---|---|---|
| SB121 | GCATCGTCTCATCGGTCTCAAACGATACCAGTTACAATTAGTATTACAAT | MET3p fw (type 3) |
| SB122 | ATGCCGTCTCAGGTCTCACATATTGTTAGGTGTTTCTTTTCT | MET3p rv (type 3) |
| SB124 | GCATCGTCTCATCGGTCTCAAACGTCTGGAACTTCTGGGAA | MT-Ip fw (type 3) |
| SB125 | ATGCCGTCTCAGGTCTCACATATGTGTTTGTTTTTGTATGTG | MT-Ip rv (type 3) |
| SB156 | GATTCTGTGGATAACCG | YTK01 entry vector seq, forward |
| SB157 | GTTCAGAACGCTCGG | YTK01 entry vector seq, reverse |
| SB226 | GTAGTGACTAAGGTTGGCC | For sequencing of Venus assemblies |
| SB202 | CCCCATCTTCGTATCTTG | For sequencing of mRuby2 assemblies |
| SB322 | GCATCGTCTCATCGGTCTCATACAAGTGCAAAGGCATTTCC | Trp1_TU_part 1 fw |
| SB323 | ATGCCGTCTCAGACTAAGTAAGCTTCAACGGATTCC | Trp1_TU_part 1 rv |
| SB324 | GCATCGTCTCAAGTCTCTTTGTTCGAAGACACC | Trp1_TU_part 2 fw |
| SB325 | ATGCCGTCTCAATACGTCAACACCAATTACACCAG | Trp1_TU_part 2 rv |
| SB326 | GCATCGTCTCAGTATCAGGTGGTGTAGAGACTGACG | Trp1_TU_part 3 fw |
| SB327 | ATGCCGTCTCAGGTCTCAACTCGACTAAATTTGGTGAAGATACTTCTG | Trp1_TU_part 3 rv |
| RP34 | AACCTCATCGCACAGCTC | For sequencing of pCgTK12 |
| RP36 | GCAAGAGGATTTCTAC | For sequencing of pCgTK12 |
| RP39 | GGCTGTCAAGAAAACTG | For sequencing of pCgTK12 |
| RP42 | CTCCACTTTGGCCAC | For sequencing of pCgTK12 |
| RP43 | GACAAGCTGATCGCAC | For sequencing of pCgTK12 |

| | | |
|---|---|---|
| RP47 | TGATAGATCCAGTAATGACC | For sequencing of pCgTK12 |
| *C. glabrata* Actin fw | TTACCGCTTTGGCTCCATCTTC | qPCR primer |
| *C. glabrata* Actin rev | TGTGGTGAACAATGGATGGACC | qPCR primer |
| *C. glabrata* Venus fw | ACCATGGGTAATACCAGCAGCA | qPCR primer |
| *C. glabrata* Venus rev | GGTGGTGTTCAATTAGCTGACCA | qPCR primer |

**Supplementary Table 5**. gRNA sequences and targeted promoters. For the target promoters 350 bp upstream of the start codon are given. gRNA targeting sites and PAM sequences are underlined and bold respectively. Note: some gRNAs target the reverse strand. The 5' UTR was extracted from the Candida genome database[4] and is given in green.

| **gRNA** | **Sequence** |
|---|---|
| URA3p #1 | atatacatgttagttaatat |
| URA3p #2 | aggaaatatgatccccttta |
| URA3p #3 | gcaagttcttgtaacttgta |
| URA3p #4 | gcatcatcccagtagattag |
| URA3p #5 | ttactatacaattccttaaa |
| ALG3p #1 | attaggactttattcggtag |
| ALG3p #2 | attttcgaaaccaaattatg |
| ALG3p #3 | cattacacaaaaaagtaacc |
| ALG3p #4 | gtcgttcattgcttatttta |
| **Promoter** | **Sequence** |
| URA3p | tttctatatacatgttatcagtcaaaccatacccataagaaaggtcaaggtaaatggtctacgtatgtttctgtgtaaaatcggattgttccgaaaatgactgctgataattgcaattttcttcatcatcgaaggactcttctcttaacggaattagctgaatagtcatctcatcg**cca**atattaactaacatgtatataaggttactatacaatt**cct**taaa**ggg**gatcatattt**cct**tacaagttacaagaacttgccagttctcgagaaccaattgcatcatcccagtagattag**tgg**atactccagctcaagactgaaaaaaaaaccatcaagagtacacttacat |
| ALG3p | gcaagacattaggactttattcggtag**agg**gcgagccactcgggtggcaagaacatgttaataatataattagtttcttagttaaaactgaaagtagctttacattacaattattttcgaaaccaaattatg**tgg**atgtcaggacttaaactacatgttagtacaaatatatctctgtgatgcatttac**ccg**gttactttttgtgtaatgcggtcgaatctagtataacttaaaa**cct**taaaataagcaatgaacgacaattgtaagtattgtgaattccattcatgagtttttaaataattgttctaaaaatataacaaaatataatacaagcattactggacattcg |

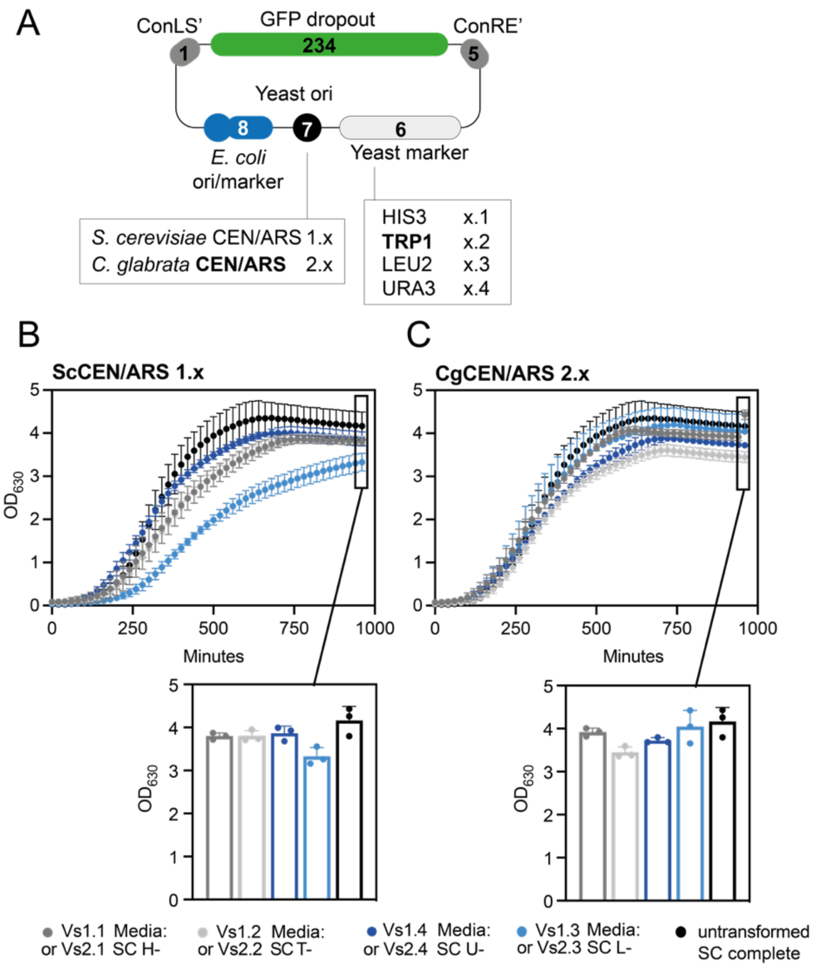


**Supplementary Figure 1. Growth of *C. glabrata* transformed with empty vectors. A.** Outline of the vector set featuring four auxotrophic markers and two origins of replication. **B.** Growth of *C. glabrata* transformed with ScCEN/ARS-vectors. Full growth curves and final $OD_{630}$ is given. **C.** Growth of *C. glabrata* transformed with CgCEN/ARS-vectors. Experiments were run in biological triplicates (three transformants) and error bars represent the standard deviation.

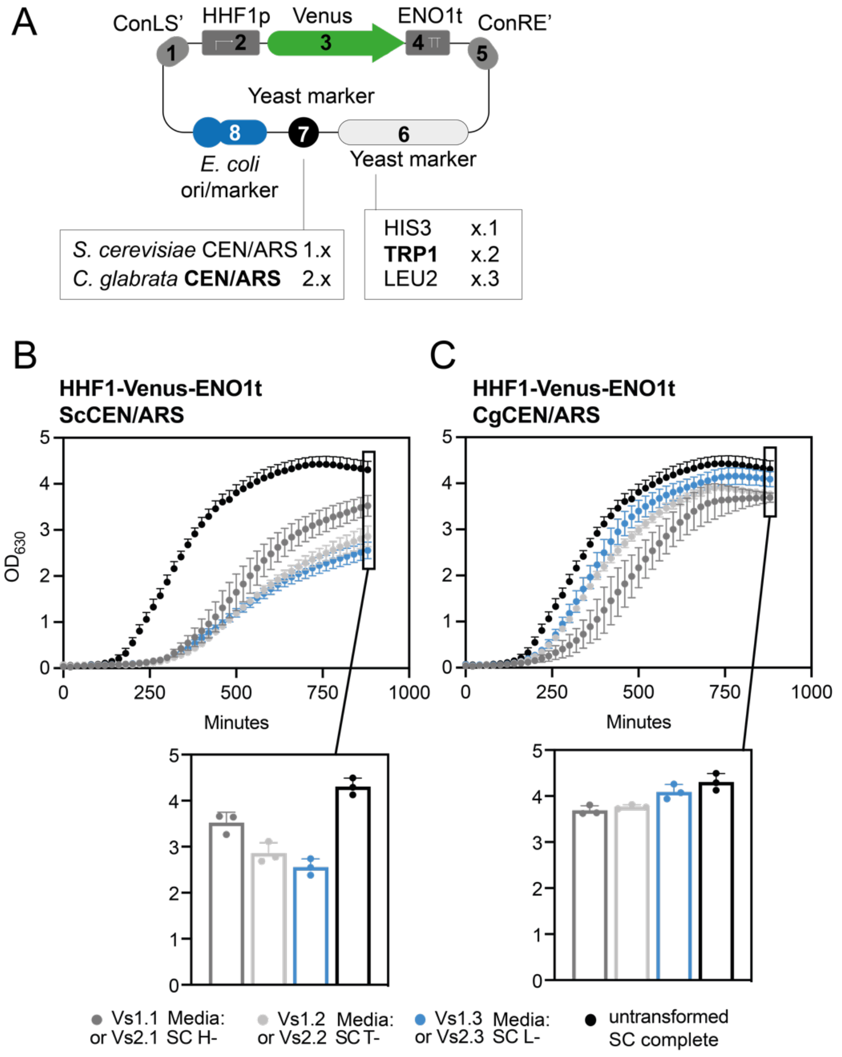


**Supplementary Figure 2. Growth of *C. glabrata* transformed with Venus-expression vectors. A.** Outline of the vector set featuring four auxotrophic markers and two origins of replication. **B.** Growth of *C. glabrata* transformed with ScCEN/ARS-vectors. Full growth curves and final $OD_{630}$ is given. **C.** Growth of *C. glabrata* transformed with CgCEN/ARS-vectors. Experiments were run in biological triplicates (three transformants) and error bars represent the standard deviation.

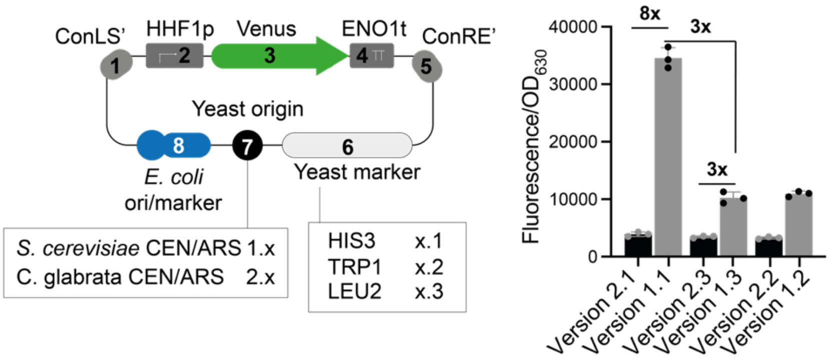


**Supplementary Figure 3.** Differences in expression level of Venus from the various vectors. **A.** Overview of used vectors. **B:** Difference in expression, numbers indicate fold difference between the indicated vectors. Experiments were run in biological triplicates (three transformants) and error bars represent the standard deviation.

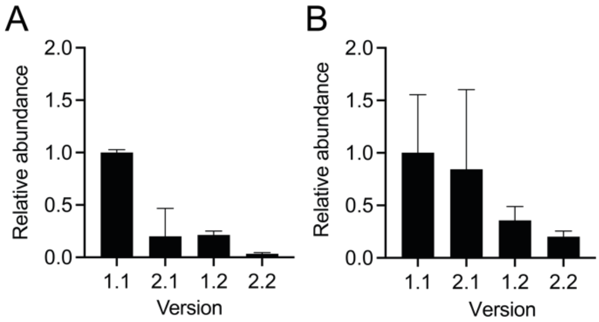


**Supplementary Figure 4.** qPCR analysis of relative cellular abundance of various vectors. **A:** qPCR on plasmid extracts **B:** qPCR based on total DNA extract. In this case plasmid copy number was normalized to actin mRNA abundance (see Methods section for both protocols).

In both (A and B), the abundance of the highest vector was set to 1 and the abundance of the other vectors is given relative to this number. Samples were measured in biological triplicates and each replicate was measured again in technical triplicate. Error bars represent standard deviation.

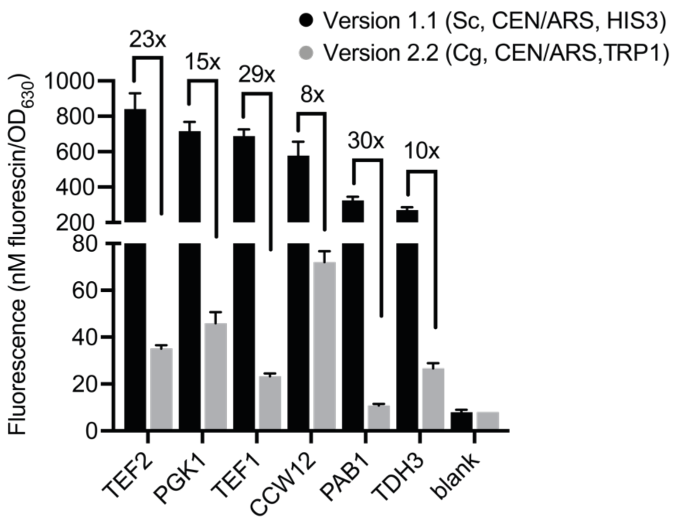


**Supplementary Figure 5.** Fold-difference in expression across various promoters when comparing the vector versions 1.1 and version 2.2. The same data as generated for Figure 2A and B were used. Experiments were run in biological triplicates (three transformants) and error bars represent the standard deviation.

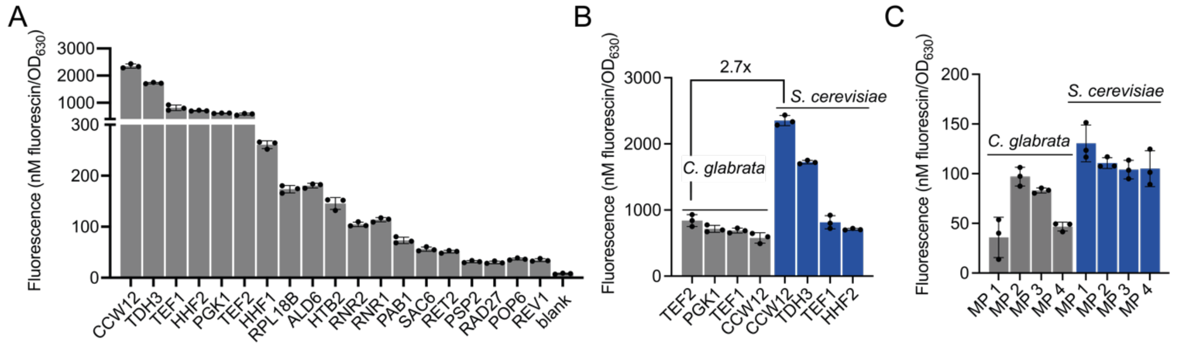


**Supplementary Figure 6.** Expression levels of YTK promoters and minimal promoters in *S. cerevisiae.* **A:** Performance of the 19 YTK promoters in *C. glabrata* using green fluorescence of the Venus protein as readout; version 1.1. **B:** The strongest promoter in *C. glabrata* (TEF2p) showed 2.7-fold lower expression than the strongest promoter in *S. cerevisiae* (CCW12p). **C:** Comparison of the expression levels resulting from the Minimal Promoters MP1-4 in *C. glabrata* and *S. cerevisiae***.** Experiments were run in biological triplicates (three transformants) and error bars represent the standard deviation.

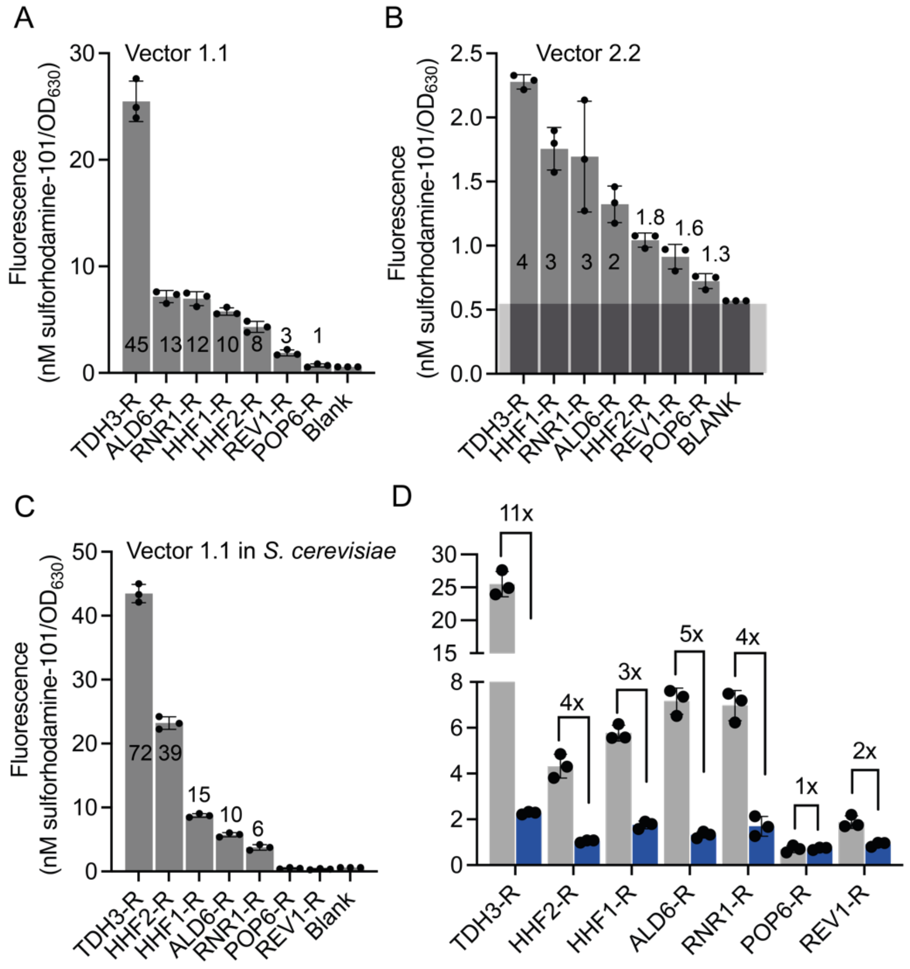


**Supplementary Figure 7. Performance of seven YTK promoters in *C. glabrata and S. cerevisiae* using red fluorescence (mRuby2) as readout. A and B:** Performance of the seven YTK promoters in *C. glabrata* using two different vectors; version 1.1 (A) and 2.2 (B). The numbers indicate the fold-change in fluorescence over background, where background is defined as the autofluorescence of *C. glabrata* cells not carrying a plasmid. Note: The Y-axis shows a different scale. **C:** Performance of the same seven YTK promoters in *S. cerevisiae* **D:** Fold difference in expression between vector version 1.1. and 2.2 in *C. glabrata*. Data from panels A and B were used. All experiments were run in biological triplicates (three transformants) and error bars represent the standard deviation.

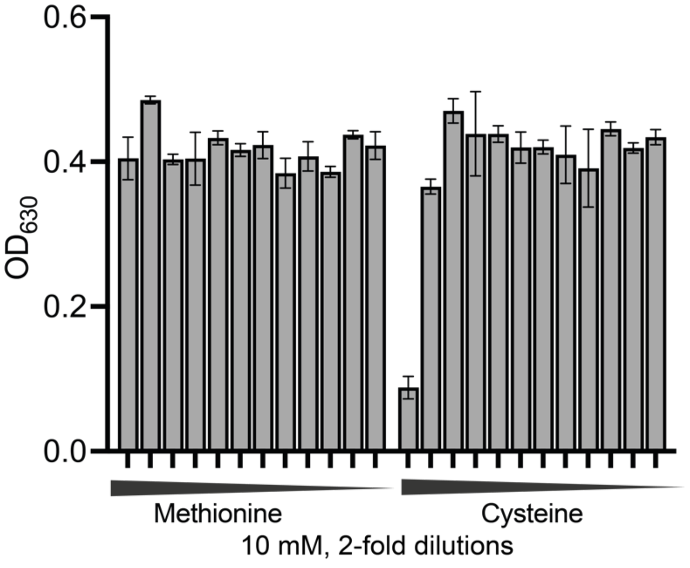


**Supplementary Figure 8.** Growth of *C. glabrata* in the presence of up to 10 mM methionine and cysteine. The final $OD_{630}$ after 24 hours of growth is shown. Experiments were run in biological triplicates (three transformants) and error bars represent the standard deviation.

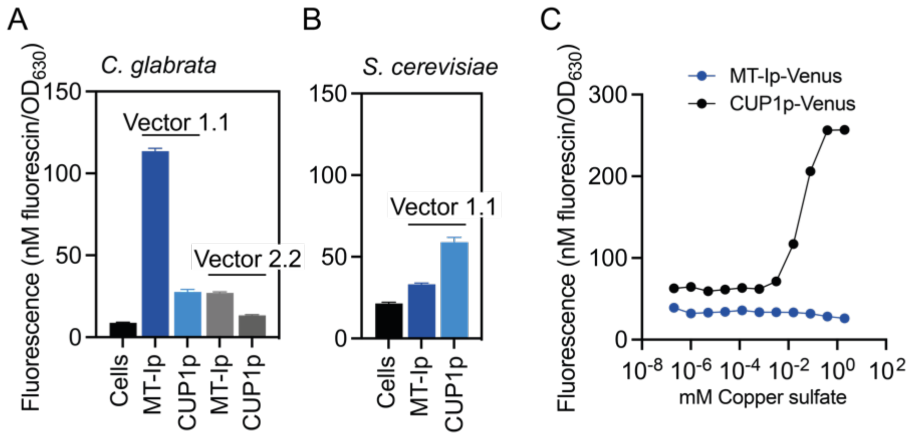


**Supplementary Figure 9. A-B.** Basal expression from the three copper inducible promoters when cloned into vector version 1.1 or 2.2 in *C. glabrata* **(A)** and *S. cerevisiae* **(B)**.
**C:** Performance of the two copper inducible promoters in *S. cerevisiae*. Experiments were run in biological triplicates (three transformants) and error bars represent the standard deviation.

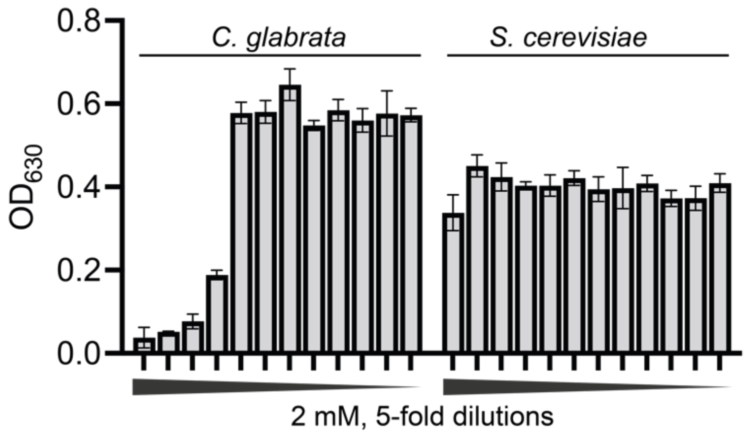


**Supplementary Figure 10.** Growth of *C. glabrata* and *S. cerevisiae* in the presence of up to 2 mM copper (II) sulfate. The final $OD_{630}$ after 24 hours of growth is shown. Experiments were run in biological triplicates (three transformants) and error bars represent the standard deviation.

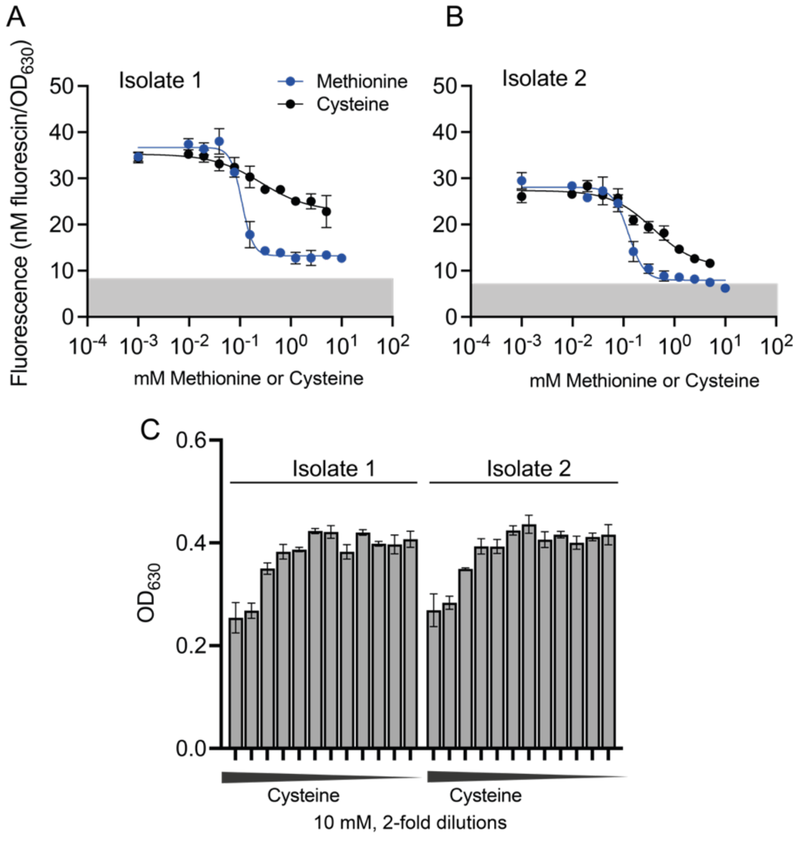


**Supplementary Figure 11. Performance of the methionine/cysteine repressible promoters in clinical isolates of *C. glabrata*. A-B:** Performance of MET3 promoters in a vector version with an ScCEN/ARS origin and a nourseothricin resistance cassette in isolate 1 (A) and 2 (B) when grown in the presence of decreasing concentrations of methionine or cysteine (starting from 10 mM methionine and 5 mM cysteine, two-fold dilutions were measured). **C:** Growth in the presence of cysteine. 10 mM cysteine led to partial growth inhibition. Experiments were run in biological triplicates (three transformants) and error bars represent the standard deviation.

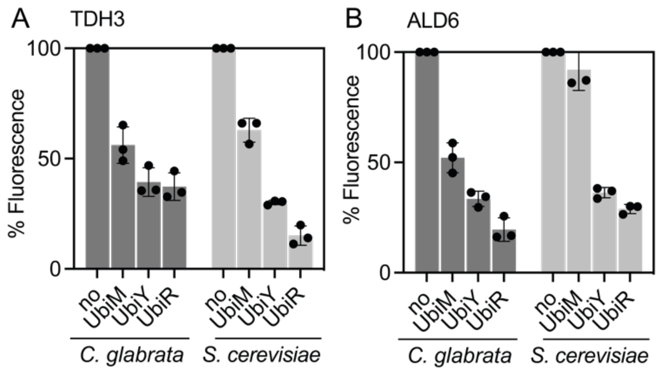


**Supplementary Figure 12.** Performance of the protein degradation tags Ubi-M, Ubi-Y, and Ubi-R in *C. glabrata* in comparison to *S. cerevisiae.* Experiments were performed with two promoters (*TDH3*, **A** and *ALD6*, **B**) in vector version 1.1. Experiments were run in biological triplicates (three transformants) and error bars represent the standard deviation.

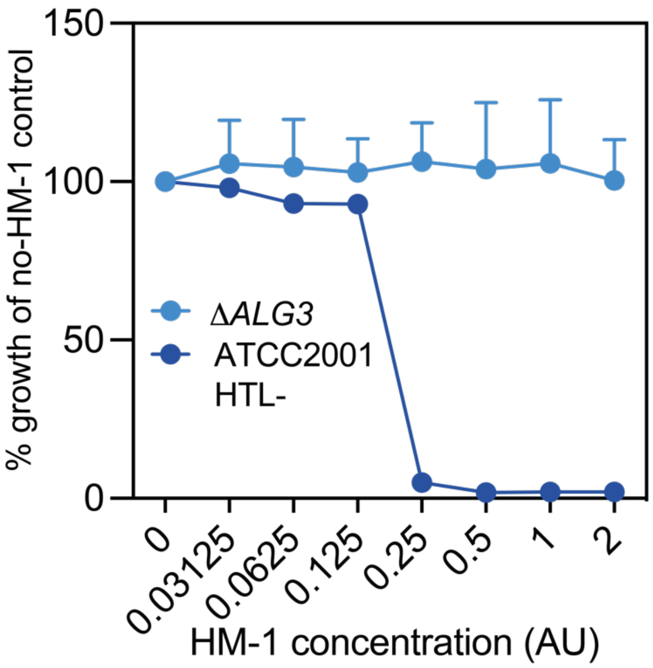


**Supplementary Figure 13.** Resistance of *C. glabrata* ATCC 2001 HTL$^-$ DeltaRHK1/ALG3 to HM-1. Cells were grown for 24 hours in the presence of increasing concentrations of HM-1. The final $OD_{630}$ was normalized to the OD of cells grown in the absence of HM-1.

1. Zordan, R. E. *et al.* Expression plasmids for use in Candida glabrata. *G3 Genes, Genomes, Genet.* **3**, 1675–1686 (2013).
2. Redden, H. & Alper, H. S. The development and characterization of synthetic minimal yeast promoters. *Nat. Commun.* **6**, 1–9 (2015).
3. Smith, J. D. *et al.* Quantitative CRISPR interference screens in yeast identify chemical-genetic interactions and new rules for guide RNA design. *Genome Biol.* **17**, (2016).
4. Skrzypek, M. S. *et al.* The Candida Genome Database (CGD): incorporation of Assembly 22, systematic identifiers and visualization of high throughput sequencing data. *Nucleic Acids Res.* **45**, D592–D596 (2017).
